# Tld1 is a novel regulator of triglyceride lipolysis that demarcates a lipid droplet subpopulation

**DOI:** 10.1101/2023.03.07.531595

**Authors:** Natalie Ortiz Speer, R. Jay Braun, Emma Grace Reynolds, Alicja Brudnicka, Jessica M.J. Swanson, W. Mike Henne

## Abstract

Cells store lipids in the form of triglyceride (TG) and sterol-ester (SE) in lipid droplets (LDs). Distinct pools of LDs exist, but a pervasive question is how proteins localize to and convey functions to LD subsets. Here, we show the yeast protein YDR275W/Tld1 (for TG-associated LD protein 1) localizes to a subset of TG-containing LDs, and reveal it negatively regulates lipolysis. Mechanistically, Tld1 LD targeting requires TG, and is mediated by two distinct hydrophobic regions (HRs). Molecular dynamics simulations reveal Tld1’s HRs interact with TG on LDs and adopt specific conformations on TG-rich LDs versus SE-rich LDs in yeast and human cells. Tld1-deficient yeast display no defect in LD biogenesis, but exhibit elevated TG lipolysis dependent on lipase Tgl3. Remarkably, over-expression of Tld1, but not LD protein Pln1/Pet10, promotes TG accumulation without altering SE pools. Finally, we find Tld1-deficient cells display altered LD mobilization during extended yeast starvation. We propose Tld1 senses TG-rich LDs and regulates lipolysis on LD subpopulations.

## Introduction

Lipid droplets (LDs) are fat storage organelles comprised of a neutral lipid core containing both triglycerides (TG) and sterol esters (SE) (Walther et al., 2017). Distinct from bilayer-bound organelles, LDs are surrounded by a phospholipid (PL) monolayer which is decorated with surface proteins that aid in their biogenesis and degradation (Currie et al., 2014). These cytosolic lipid reservoirs can be made or broken down in response to a variety of metabolic cues, such as nutrient deprivation or increased membrane biogenesis. Defects in lipid storage in LDs contribute to numerous metabolic disorders including obesity, cardiovascular disease, and diabetes (Welte, 2015; Gluchowski et al., 2017). Recent studies indicate that beyond their role in lipid storage, LDs also play important roles in signaling and protein homeostasis (Li et al., 2012; Bersuker et al., 2018; Schmeisser et al., 2019). Despite this, it remains unclear if distinct pools of LDs exist within cells to enable this functional diversity. Work from our group and others have shown that LDs are not homogenous within the context of a single cell, but exist in a variety of subpopulations that contain distinct proteomes and/or morphologies (Zhang et al., 2016; Eisenberg-Bord et al., 2018; Teixeira et al., 2018; Schott et al., 2019; Ugrankar et al., 2019). Although LDs exhibit these unique features, little is currently known regarding how such differences dictate LD function. LD subpopulations are of particular interest to the field of metabolism as there is mounting evidence that different LD pools play roles in maintaining metabolic homeostasis in response to various nutrient states (Hariri et al., 2018; Eisenberg-Bord et al., 2018; Teixeira et al., 2018). For example, large and small LD pools observed in human hepatocytes are mobilized by mechanistically distinct pathways during starvation (Schott et al., 2019). Similarly, *Drosophila* fat body cells contain two subpopulations of LDs that are differentially maintained by extracellular and *de novo* synthesis of lipids (Ugrankar et al., 2019).

LD turnover primarily occurs through a highly conserved process known as lipolysis. This catabolic process involves the targeting of cytoplasmic lipases to LDs where they hydrolyze TG and SE to base components. TG breakdown via lipolysis is necessary for maintaining lipid homeostasis, sustaining membrane biosynthesis, and promoting cellular division across multiple species (Duncan et al., 2007; Schmidt et al., 2014; Heier and Kühnlein, 2018). However, the underlying mechanisms for regulation of TG lipolysis in budding yeast are poorly understood. Yeast contain three LD-resident and paralogous TG lipases: Tgl3, Tgl4, and Tgl5 (Athenstaedt et al., 1999; Athenstaedt and Daum, 2003, 2005; Kurat et al., 2006). Although Tgl4 has been shown to be the functional ortholog of the mammalian TG lipase, ATGL, in yeast, it is in fact Tgl3 that performs the bulk of the lipolytic activity *in vivo* as it can hydrolyze TG species of variable fatty acid chain length (Athenstaedt and Daum, 2003, 2005; Kurat et al., 2006). The regulation of Tgl3-mediated TG lipolysis is poorly understood. Previous studies provide some insight by demonstrating that in the absence of either TG or LDs as whole, Tgl3 activity, targeting, and stability is negatively impacted, a common trait for many resident LD proteins (Schmidt et al., 2013; Koch et al., 2014). In LD-null yeast, Tgl3 is re-targeted to the ER where it loses its lipolytic activity and is rapidly degraded (Schmidt et al., 2013). In spite of this information, specific regulators of Tgl3 TG lipase activity remain unidentified. Whether specific LD subsets are preferentially mobilized during metabolic cues is also underexplored.

Here, we deorphanize the LD protein YDR275W/Bsc2, a poorly understood LD targeting protein in yeast. We find it decorates LD subsets and acts as a negative regulator of TG lipolysis. Given these properties we propose to name it Tld1 (for <u>T</u>G-associated <u>LD</u> protein <u>1</u>). We find Tld1 LD targeting is dependent on the presence of TG, as Tld1 fails to stably localize to SE-LDs in yeast or human cells. Structure-function analysis reveals the N-terminal half of Tld1, containing distinct hydrophobic domains, is necessary for stable LD association. This is supported by molecular dynamics (MD) simulations that demonstrate Tld1 adopts a distinct conformational ensemble on TG-rich LDs and interacts directly with TG in addition to LD monolayer PLs. Physiologically, loss of Tld1 causes significantly decreased TG levels during yeast LOG phase growth. This decrease is not due to reduced TG synthesis, but rather from upregulated Tgl3-mediated TG lipolysis. Conversely, Tld1 over-expression promotes TG accumulation and LD enlargement, but does not alter SE levels. Thus, we propose that Tld1 demarcates a LD subpopulation where it locally inhibits Tgl3-dependent TG lipolysis.

## Results

### Tld1 localizes to a LD subset and requires TG for LD targeting

To dissect how proteins target to specific lipid droplets (LD) subpopulations, we used a candidate-based approach to image GFP-tagged proteins annotated to localize to LDs in the budding yeast *Saccharomyces cerevisiae*. We manually imaged yeast expressing chromosomally GFP-tagged LD proteins and co-expressing the canonical LD protein Erg6-mRuby, a previously established LD marker that decorates all LDs (Müllner et al., 2004). Candidate-based imaging revealed that GFP-tagged YDR275W/Bsc2 (as discussed below, here on referred to as Tld1 to avoid confusion with Bscl2, another name for seipin), was detected on only a subset of Erg6-mRuby labeled LDs in yeast growing at logarithmic (LOG) phase (**Fig 1A**). Tld1 is annotated as an LD protein, but its function remains uncharacterized. Similar to its colocalization with Erg6-mRuby, LOG-phase yeast expressing Tld1-GFP and stained with the general LD dye monodansylpentane (MDH) also showed partial MDH and Tld1-GFP co-localization (**Fig 1C**). Consistent with this, previous work also determined that Tld1 was among a few LD proteins detected on LD subsets in budding yeast (Eisenberg-Bord et al., 2018; Teixeira et al., 2018). To determine whether Tld1-GFP decorated a LD subset in yeast in different growth phases, we also imaged yeast grown into stationary (STAT) phase, when cell growth slows and LD lipid storage is elevated. STAT phase yeast also exhibited detectable Tld1-GFP on LDs, but this Tld1-GFP signal colocalized closely with Erg6-mRuby (**Fig 1A**). Quantification of this Tld1-GFP/Erg6-mRuby colocalization in LOG and STAT phases revealed that in LOG phase, only ∼40% of Erg6-mRuby LDs also exhibited detectable Tld1-GFP (**Fig 1B**). In STAT phase this detectable co-localization increased to ∼70%, suggesting Tld1-GFP and Erg6-mRuby colocalization increased in STAT phase. We also monitored Tld1 expression in LOG and STAT phase yeast by Western blotting for endogenous GFP-tagged Tld1-GFP strains using an anti-GFP antibody. Consistent with the fluorescence imaging, Tld1-GFP appeared slightly more abundant in STAT phase yeast compared to LOG phase, consistent with the model that Tld1-GFP accumulates in STAT phase yeast (**Fig 1D).**

**Figure 1.**
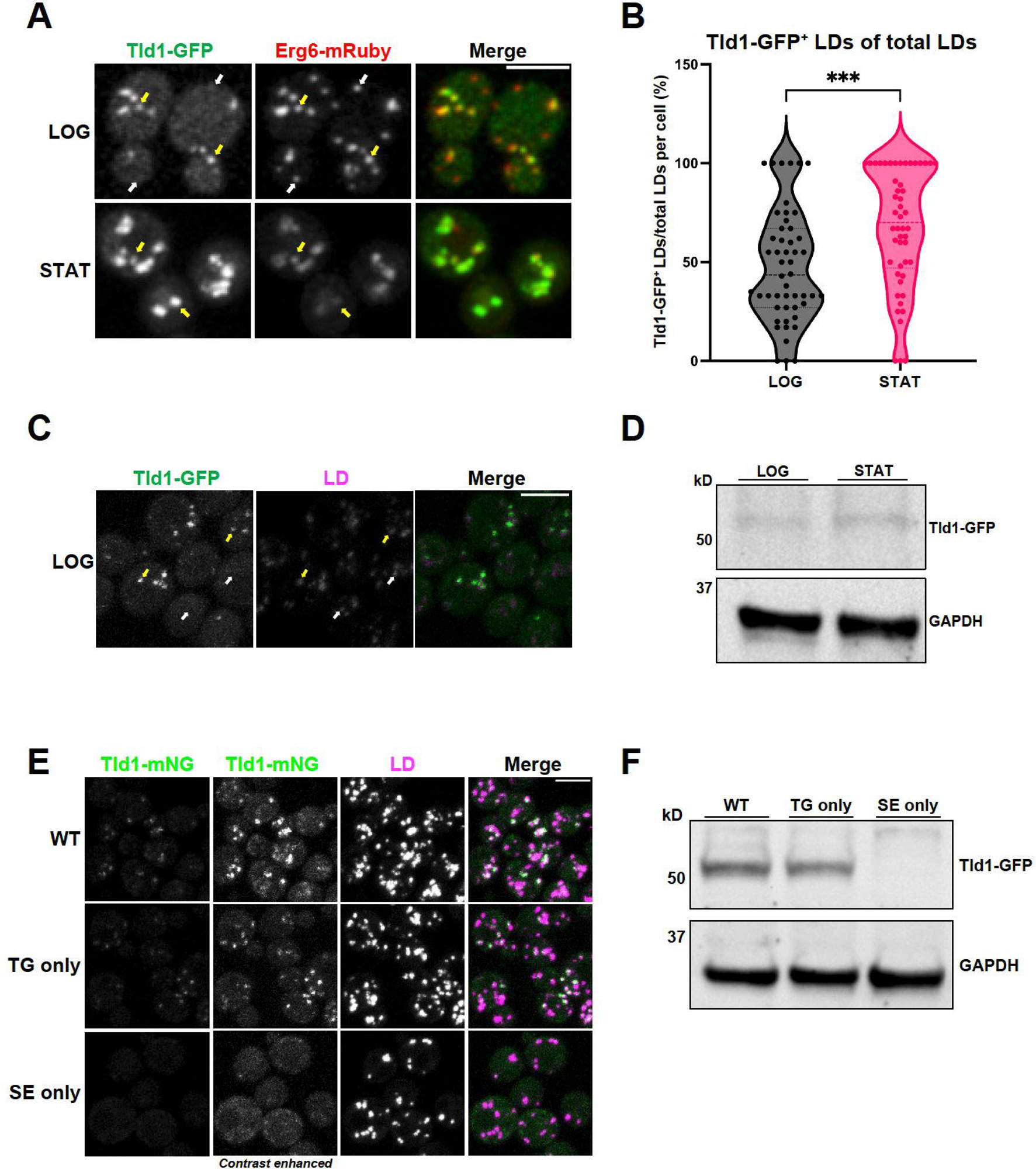
Tld1 enriches on a TG-containing LD subpopulation at logarithmic phase. **(A)** Logarithmic (LOG) and stationary (STAT) phase imaging of yeast dual-tagged for Tld1-GFP, Erg6-mRuby. Yellow arrows indicate Tld1-enriched LDs and white arrows indicate LDs where Tld1 is undetectable or absent. **(B)** Quantification of percentage of Tld1-positive (Tld1^+^) LDs out of total Erg6-mRuby LDs, per cell, at LOG and STAT phase. For both LOG and STAT samples, n = 50 cells. **(C)** Tld1-GFP expressing yeast stained with LD dye MDH and imaged at LOG phase growth. Yellow arrows are Tld1-positive LDs, white arrows denote Tld1-negative LDs. **(D)** Protein expression of Tld1-GFP in yeast grown to LOG and STAT. **(E)** Imaging of Tld1-mNeonGreen (Tld1-mNG) yeast in different neutral lipid-containing backgrounds with MDH-stained LDs at LOG phase. TG = Triglyceride, SE = Sterol Ester. Far left column represents non-contrast adjusted images for Tld1-mNG. **(F)** Protein expression levels of Tld1-GFP in WT, TG only, and SE only yeast. Statistics represent Unpaired t test with Welch’s correction. ***, P < 0.001. Scale bars, 5µm.

Recent work indicates that LD neutral lipid composition can influence protein targeting to the LD surface (Thiam and Beller, 2017; Chorlay and Thiam, 2020; Caillon et al., 2020; Dhiman et al., 2020). Since yeast LDs contain TG and SE, we next dissected whether loss of either of these neutral lipids influenced Tld1-mNeonGreen (Tld1-mNG) LD localization. We generated a chromosomally-tagged Tld1-mNG yeast strain that produced only TG (TG-only) by deleting the genes encoding the two SE-generating enzymes Are1 and Are2, and a strain producing only SEs (SE-only) by deleting the TG-synthesis enzymes Dga1 and Lro1 (Sandager et al., 2002; Sorger et al., 2004). Imaging revealed that whereas the wildtype (WT) and TG-only yeast exhibited Tld1-mNG that co-localized with a subset of LDs, the SE-only yeast contained very dim Tld1-mNG signal that was nearly undetectable on LDs (**Fig 1E**). To determine whether the dim Tld1-GFP signal in SE-only yeast was due to lower Tld1-GFP protein abundance, we Western blotted for endogenous Tld1-GFP in wildtype, TG-only, and SE-only yeast. Indeed, SE-only yeast displayed a near complete loss of Tld1-GFP compared to the other yeast stains (**Fig 1F**). Collectively, this suggests that TG is necessary for Tld1 LD targeting and to maintain Tld1 protein levels *in vivo*. Due to its apparent targeting preference for TG-containing LDs we propose re-naming this protein Tld1 for <u>T</u>G-associated <u>LD</u> protein 1. This name also avoids confusion between Bsc2 and Bscl2, which is another name for the LD associated protein seipin.

### The Tld1 N-terminal hydrophobic regions mediate LD targeting

Proteins can target to LDs through amphipathic or hydrophobic motifs that interact with or insert into the LD PL monolayer (Bacle et al., 2017; Prévost et al., 2018; Chorlay and Thiam, 2020; Chorlay et al., 2021). To mechanistically dissect how Tld1 targets to LDs, we examined its hydrophobicity using Phobius (Käll et al., 2004) (**Fig 2A**). The hydrophobicity plot predicted two hydrophobic regions in the N-terminal half of Tld1, which we denote as Hydrophobic Region 1 (HR1) and Hydrophobic Region (HR2). Tld1 also contains a predicted Low Complexity Region (LCR) directly downstream of these HRs. We hypothesized that Tld1 targets to LDs through the action of either HR1, HR2, or both regions. To test this, we generated seven mNG-tagged fragments of Tld1, and over-expressed them in yeast stained for LDs in LOG phase growth (**Fig 2B**). We also conducted Western blotting to access the expression levels of these constructs (**Supplemental Figure 1A**). Interestingly, full length Tld1 (Tld1^FL^) targeted both LDs and the endoplasmic reticulum (ER) when over-expressed. Similarly, a truncated fragment removing the LCR (Tld1^N-HR1+HR2^) also showed this LD and ER dual-targeting, as did a smaller fragment only containing the HR1 and HR2 regions (Tld1^HR1+HR2^), suggesting the LCR and N-terminal region (Tld1^N^) preceding HR1 are not necessary for this LD/ER localization. We noted that the small Tld1^N^ construct generally distributed in the cytoplasm and did not enrich on LDs.

**Figure 2.**
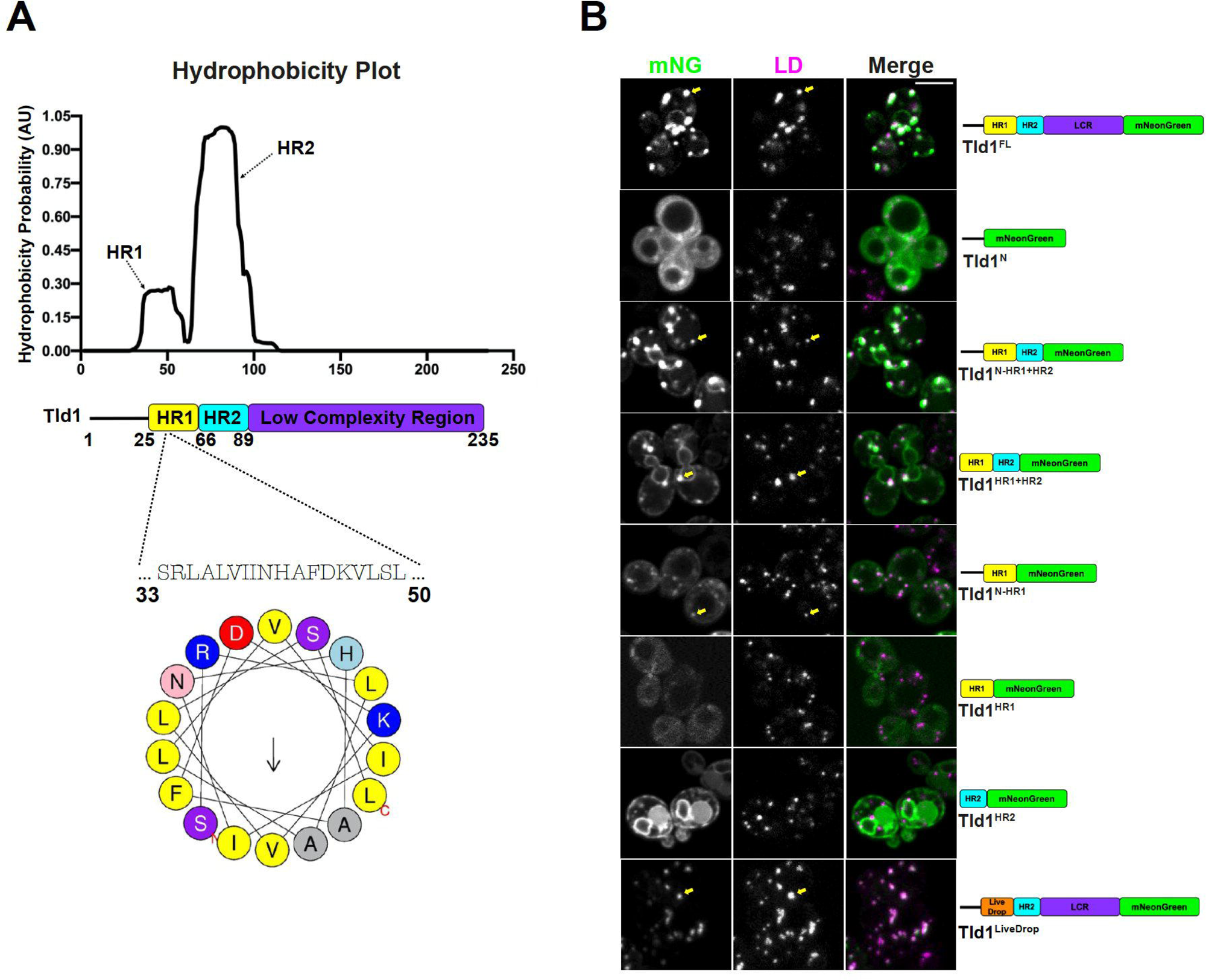
Hydrophobic Region 1 (HR1) of Tld1 is responsible for LD-targeting. **(A)** Hydrophobicity plot generated by Phobius online transmembrane topology and signal peptide predictor (top), paired with a schematic of Tld1 protein architecture (middle), and a helical wheel of the Tld1 amphipathic segment in HR1 (bottom). HR1 = Hydrophobic Region 1, HR2 = Hydrophobic Region 2. **(B)** Log phase imaging of yeast over-expressing various Tld1-mNG truncations with LDs stained with MDH. Images are midplane sections. Yellow arrows indicate LD-targeting. LCR = Low Complexity Region.

Next we dissected how HR1 and HR2 influence Tld1 localization to the ER network and LDs. A construct encoding only the N-terminal region and HR1 (Tld1^N-HR1^) localized to LDs, suggesting HR1 may be sufficient for LD targeting (**Fig 2B**). In support of this, amino acid and secondary structure analysis of HR1 indicates it forms a predicted alpha-helical fold, with several hydrophobic amino acids on one face, commonly observed in LD targeting motifs (**Fig 2A**). A smaller construct retaining HR1 without the preceding N-terminal region (Tld1^HR1^) failed to express well in yeast (**Supplemental Figure 1A**), suggesting the initial N-terminal region may be necessary for HR1 stability. Surprisingly, a construct encoding only HR2 (Tld1^HR2^) localized primarily to the ER network surrounding the nucleus and peripheral ER (**Fig 2B**). No detectable LD localization was observed for Tld1^HR2^, indicating HR1 was necessary for LD targeting. Since HR1 appeared to be required the Tld1 LD interaction, we generated a chimeric Tld1 construct where we replaced HR1 with LiveDrop (Tld1^LiveDrop^), a known LD targeting module derived from the LD targeting motif of *Drosophila* GPAT4 (Wilfling et al., 2013; Wang et al., 2016). Indeed, Tld1^LiveDrop^ targeted to LDs and appeared similar to Tld1^FL^, suggesting LiveDrop could replace HR1 for organelle targeting (**Fig 2B**).

Tld1 LD targeting could, in principle, be due to direct protein insertion or interaction with the LD surface, or through Tld1 binding to another yeast LD protein. To delineate these possibilities, we expressed full length yeast Tld1-EGFP in human U2-OS cells treated with oleic acid (OA) to induce LD biogenesis. Tld1-GFP decorated the surfaces of LDs in U2-OS cells, suggesting it was able to localize to the LD surface independent of other yeast proteins (**Fig 3A**). Similar to yeast, Tld1-EGFP was only detected on a subset of LDs in human cells. Next, we expressed EGFP-tagged Tld1^HR1^, Tld1^HR2^, and Tld1^HR1+HR2^ fragments in OA-treated U2-OS cells. Similar to their localization patterns in yeast, Tld1^HR1+HR2^-EGFP localized to both the ER network and LD surfaces. Tld1^HR1^-EGFP decorated LD surfaces as well as localized in a diffuse pattern in the cytoplasm, again suggesting HR1 was sufficient for LD targeting. Tld1^HR2^-EGFP targeted exclusively to the ER network and nuclear envelope with no detectable enrichment on LD surfaces, similar to its localization when expressed in yeast. Collectively, this supports a model where Tld1 can target to both the ER and LD, and interacts directly with the LD surface independent of other yeast proteins. It also suggests that HR1 is necessary and sufficient for LD targeting, whereas HR2 alone favors an ER localization. We speculate that the presence of HR2 may promote HR1 LD targeting by providing local enrichment of Tld1 on the ER surface near LD budding sites.

**Figure 3.**
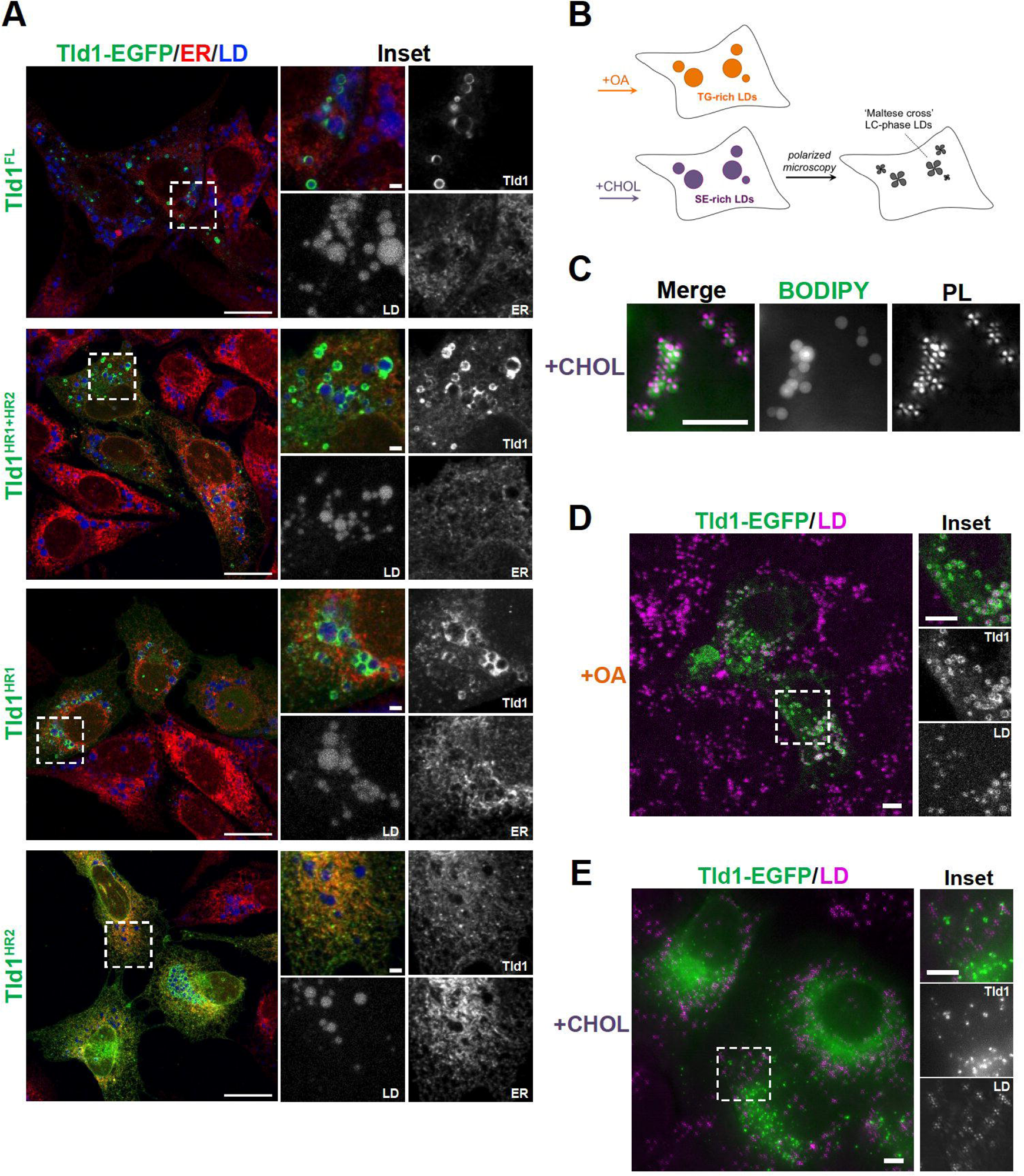
Tld1 targets to and prefers TG LDs in mammalian cells. **(A)** Confocal imaging of U2-OS cells transiently overexpressing either Tld1 or Tld1 fragments, all tagged with EGFP (Tld1-EGFP) and treated with oleic acid (OA), overnight to induce LD formation. Cells were coIF stained α-HSP90B1 (ER, red) and LDs stained with MDH (blue) and imaged with confocal microscope. Scale bar, 20 μm. Inset scale bar, 5µm. **(B)** Schematic of oleic acid (OA) and cholesterol (CHOL) treated HeLa cells to generate TG-rich or SE-rich LDs. Maltese crosses bottom right cell represent liquid crystalline (LC) phase, SE-rich LDs, visible by polarized light microscopy. **(C)** Polarized light (PL) microscopy of HeLa cells treated with cholesterol to form SE-rich. LDs were stained with BODIPY 493/503 (green) and visualized by polarized light (magenta crosses). Scale bar, 5µm. **(D)** Confocal imaging of Tld1-EGFP expressing HeLa cells, treated with OA to form TG-rich LDs. LDs were stained with MDH (magenta). Scale bars, 10µm. **(E)** Confocal imaging of Tld1-EGFP expressing HeLa cells, treated with CHOL to form SE-rich LDs. LDs were visualized with polarized light (magenta). Scale bars, 10µm.

### Tld1 displays binding preference for TG-rich LDs in human cells

Since Tld1 appeared to require TG for LD targeting in yeast, we also monitored how LD neutral lipid composition influenced Tld1-EGFP LD localization in human cells. We treated HeLa cells with either BSA-conjugated OA or cyclodextrin-conjugated cholesterol to generate TG-rich or SE-rich LDs, respectively, using a protocol previously used in our lab (Rogers et al., 2022) (**Fig 3B**). Interestingly, we could monitor SE-rich and TG-rich LDs by staining them with BODIPY, or with polarized light microscopy. This is because the hydrophobic core of SE-rich LDs forms a smectic liquid-crystalline (LC) phase that diffracts polarized light and exhibits a “Maltese cross” pattern in polarized light microscopy (**Fig 3C**). TG-rich LDs lacking this LC phase do not exhibit this Maltese cross (Rogers, et al 2022).

Equipped with this method, we transfected cells with Tld1-EGFP and monitored its localization. As expected, Tld1-EGFP localized strongly to the surfaces of TG-rich LDs stained with MDH (**Fig 3D**). In contrast, Tld1-EGFP did not detectably enrich on the surfaces of Maltese-cross SE-rich LDs in cholesterol-treated cells (**Fig 3E**). However, we detected numerous small Tld1-EGFP foci in cholesterol-treated cells, but these did not colocalize with the Maltese crosses of SE-rich LDs. We speculate that these foci may represent protein aggregates or potentially sites of TG accumulation that Tld1 associates with, but this will require additional study. Collectively, this suggests that, similar to yeast, Tld1-EGFP exhibits preference for binding TG-rich LDs.

#### Molecular dynamics simulations suggest Tld1 HRs adopt specific conformations on TG-rich LDs

We next investigated how Tld1 interacted with LDs and showed preference for TG-containing LDs. Molecular dynamics (MD) simulations were conducted with Tld1^N-HR1+HR2^ (residues 1-100) interacting with a TG-rich LD (100% TG), a SE-rich LD (90:10 ratio of cholesteryl oleate (CHYO) to TG), and an ER bilayer. The structure of Tld1^N-HR1+HR2^ was first predicted with RoseTTAFold (Baek et al., 2021) and AlphaFold2 (Jumper et al., 2021), both of which predicted an alpha-helix for HR1, and a hairpin (helix-kink-helix) conformation for HR2. TOPCONS (Tsirigos et al., 2015) and TM AlphaFold (Dobson et al., 2023) also predicted a membrane-embedded topology for the hydrophobic HR2 sequence (**Supplemental Fig 2M**). Although they were very similar, the RoseTTAFold structure was selected for further simulations as it has been demonstrated to better predict membrane structures (Azzaz et al., 2022; Hegedűs et al., 2022). Tld1 HR2 was then embedded into each lipidic environment deep enough to enable the charged residues at the top of the hairpin (Arg61, Asp90, Asp93, Arg100) to be surface oriented (**Fig 4A, Supplemental Fig 2C**). HR1 was positioned 5 Å above each membrane to track its potential association with membrane packing defects (see Methods). Long timescale simulations were run on Anton2 provided by Pittsburg Supercomputing Center (Shaw et al., 2014), yielding 4.5 µs of simulations for the TG-rich LD and ER bilayer systems. Due to limited Anton2 time, the SE-rich LD system was run for 1 µs on EXPANSE provided by San Diego Supercomputing Center (Strande et al., 2021). The RMSDs suggest convergence for HR1 and minimal changes in all systems for HR2 after 1 µs (**Supplemental Fig 3A**). Since the N-terminus shows similar interactions for the TG-rich LD and bilayer (**Supplemental Fig 3B, 3C**), we focus below on HR1 and HR2.

**Figure 4:**
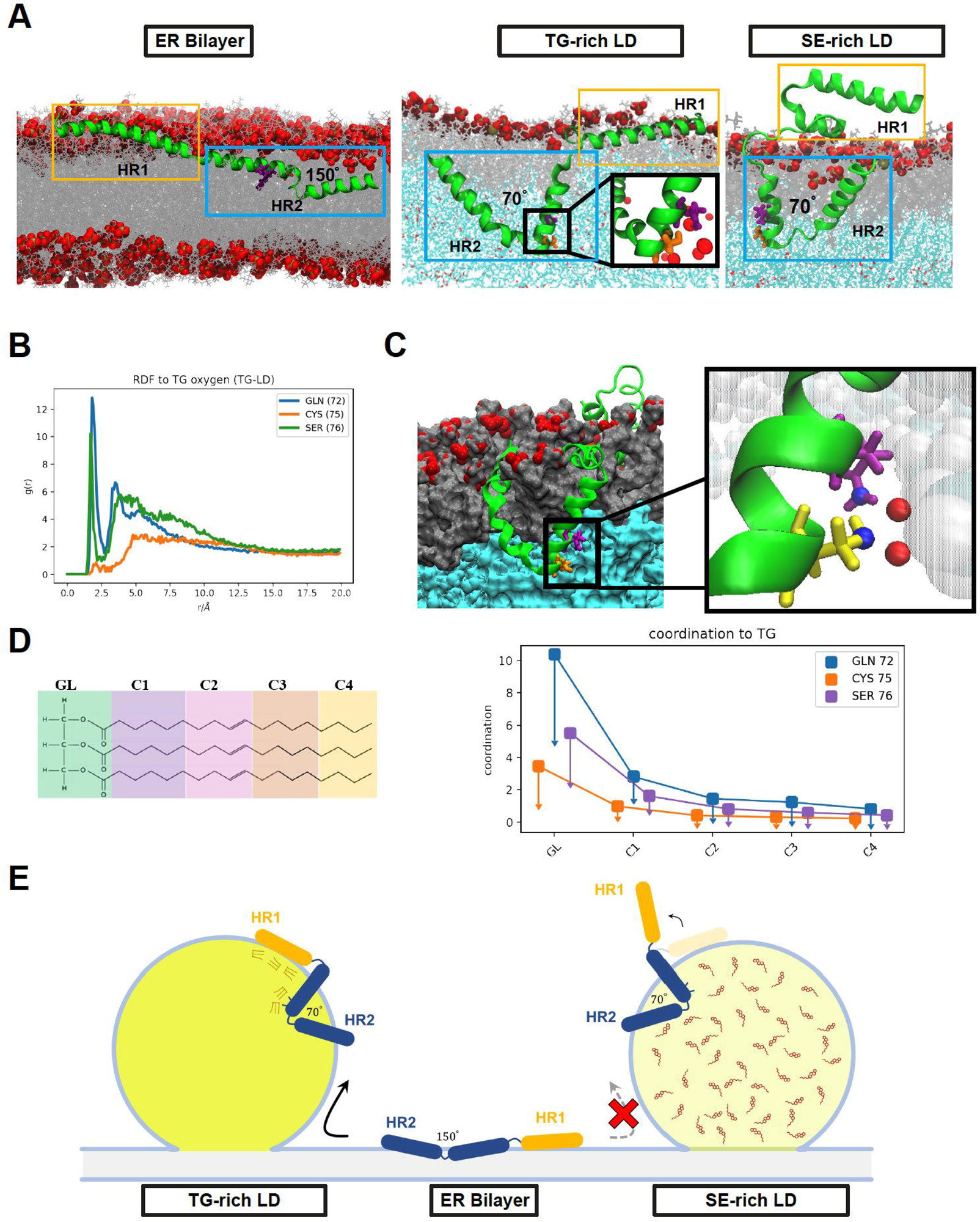
Molecular dynamics simulations indicate Tld1 adopts a unique conformational ensemble on TG-rich LDs. (**A**) After equilibration, Tld1^N-HR1+HR2^ adopts unique structures in the ER bilayer and LD monolayers. In the bilayer (left), the HR2 sequence opens to allow for polar residues in the kink to evade the unfavorable PL tail region. In a TG-rich LD (middle) these polar residues are stabilized by TG-glycerol headgroups in the LD core. In the SE-rich LD (90:10 CHYO:TG LD, right), HR2 retains the kinked conformation with the polar residues stabilized by CHYO oxygens in the core. However, the amphipathic HR1 sequence fails to associate due to significantly decreased packing defects. (**B**) Radial distribution functions of GLN, CYS, and SER in the HR2 interacting with TG glycerols. **(C)** A cross section of the LD monolayer highlights interactions between GLN72 (purple), SER76 (yellow) and TG oxygens (inset). **(D)** The coordination number between residue heavy atoms (right) and different sections of the TG molecules (left) verifies that most interactions are with the glycerol (GL) group. (**E**) Schematic of modeled Tld1^N-HR1-HR2^ adopting conformations in the TG-rich LD, ER bilayer, and SE-rich LD as in panel 4A.

Simulations revealed clear conformational changes between the LD and ER bilayer environments (**Fig 4A, Supplemental Fig 3D-F**). In both the TG-rich and SE-rich LDs, HR2 initially orients with the predicted helix-kink-helix angle of approximately 100^°^, then decreases to an angle of 70^°^ as the kink region engages with the TG core (**Supplemental Fig 2A, 2B)**. In contrast, in the ER bilayer the helix-kink-helix region opens to an average angle of 150^°^, bringing the residues in the kink region closer to the PL surface (**Fig 4A, Supplemental Fig 2A, 2B**). A central driving force for this conformational change is the stabilization of polar residues Gln72, Cys75 and Ser76 near the kink of HR2. In the LDs, these residues interact with TG glycerol groups 2.0-2.5nm below the headgroup phosphates (**Fig 4B, 4C, Supplemental Fig 2E**). In the ER bilayer, stabilization at this depth is not possible as it places the polar residues in the hydrophobic tail region of the PLs. By splaying open, the HR2 kink region rises closer to the lipid head-groups, enabling polar interactions with the PL-glycerols ∼1-1.2nm below the phosphate plane (**Supplemental Fig 2F**). Thus, HR2 obtains a more kinked conformation in the LD monolayers, but a splayed open conformation in the ER bilayer.

Focusing on HR1, the amphipathic helix embeds well in the surface packing defects, areas where hydrophobic PL acyl chains or neutral lipids are exposed to the cytosol, within the first 50 ns of the simulation (**Fig 4A, Supplemental Fig 3G**). These defects stabilize hydrophobic residues. Contact analysis, which quantifies the number of protein and membrane heavy atoms withing 3 Å, demonstrates that the hydrophobic residues along the bottom of HR1 interact with both PL and TG acyl tails (**Supplemental Fig 2H-J, 3G**), while the charged and polar residues along the top stabilize the HR1 helix via hydrogen bonds with the PL headgroups and water. Strikingly, this is not the case for the SE-rich LD. Here the HR1 helix fails to associate with the monolayer surface, and instead folds over on itself to maintain some degree of amphipathic interactions (**Fig 4A, Supplemental Fig 3F, 2K**). The reason for this is insufficient lipid packing defects in the SE-rich LD to stabilize the HR1 hydrophobic moieties. Thus, the amphipathic helix HR1 associates well with the TG-rich LD and ER bilayer, but fails to associate at all with the SE-rich LD (depicted in **Fig 4E** cartoon).

Based on these simulations, the driving force for Tld1 LD targeting is likely a combination of the Tld1 HR1 and HR2 sequences working together. Due to its drastically different confirmation on TG-rich versus SE-rich LDs, it is possible that HR1 acts as a ‘sensor’, detecting the numerous packing defects found on TG-rich LDs preferentially over SE-rich LDs and the ER bilayer. TG-rich LDs have been shown to have larger and longer-lived packing defects, with a packing defect constant of 27Å^2^, than the ER bilayer (only 16 Å^2^) (Kim et al., 2021; Braun and Swanson, 2022). This discrepancy is even more pronounced for the SE-rich LD, which has a more densely packed PL monolayer with very few packing defects, maintaining a defect constant of 14Å^2^ (Braun and Swanson, 2022). Collectively, the preferential targeting of HR1 to TG-enhanced packing defects could explain why the over-expressed Tld1^N-HR1^ fragment localizes to LDs, and also provides a molecular explanation for why Tld1 localizes to TG-rich LDs but appears significantly less detectable on SE-rich LDs *in vivo*.

The hydrophobic HR2 segment seems to embed in either the ER bilayer or LD monolayers. We hypothesize that in the absence of HR1, Tld1^HR2^ remains in the ER of both yeast and human cells either because it is thermodynamically more stable there or it is kinetically trapped in a splayed-open conformation. However, in the presence of HR1, HR2 may fold into a more stable kinked conformation in TG-rich LDs once the polar residues (Gln72, Cys75, Ser76) gain access to the glycerol groups of TG molecules in the LD core (**Fig 4A**). This is supported by the depths of Gln72, Cys75 and Ser76 in the TG-rich LD (**Supplemental Fig 2E**), as well as free energy profiles of the wildtype and mutant variants discussed below. Additionally, radial distribution functions (RDF) and coordination numbers 1s1 verify there are strong interactions between Gln72 and Ser76, especially to TG oxygens, while the hydrophobic residues surrounding these polar residues are still stabilized by PL tails (**Fig 4B-D**). In contrast, in the ER bilayer the HR2 region opens into a shallower interfacial conformation below the PL headgroups because of the high barrier for the polar residues to enter the PL tail region (**Supplemental Fig 2F, Supplemental Fig 3D**). The relative stability of these two regions is captured in the potential of mean force (PMF) profiles for amino acid permeation (**Supplemental Fig 2G**), showing that Gln72 and Ser76 are most stable slightly below the PL phosphate groups, where the polar backbone and sidechain groups can interact with the polar PL components. Pulling them into the lipid tail region is highly unfavorable. Considering Ser76 alone, moving from its interfacial position (∼1.0 nm below the phosphate plane) (**Supplemental Fig 2G, circle)** to the depth of the LD kinked position (∼2.2 nm below) (**Supplemental Fig 2G, triangle)** would cost ∼10 kcal/mol. Such high penetration barriers would explain a kinetic barrier keeping Tld1^HR2^ localized in the ER bilayer. In this case, HR2 may offer a stabilizing force once taken to the LD in the presence of HR1, which could overcome the kinetic barrier to enable HR2 to transition to a more stable LD conformation. Alternatively, Tld1^HR2^ may be more thermodynamically stable in the ER.

It is also notable that Tld1 interacts with many TG molecules in the TG-rich LD system. HR2 coordinates with the TG-glycerol backbone, and HR1 forms several contacts with TG hydrophobic tails that intercalate into the PL monolayer (**Fig 4C, 4D, Supplemental Fig 2H-J**). Thus, the LD core appears to require an abundance of TGs for optimal Tld1 interactions. The proportion of conformations with a TG molecule directly interacting with a residue captures the abundance of these interactions (**Supplemental Fig 2L**). The dominance of TG-interactions in the HR2 region demonstrates the sequence disposition to immerse itself within a TG-rich LD core. Additionally, the number of contacts between HR1 and TG-tails is a significant addition to its interactions with the PL-tails (**Supplemental Fig 2J**). Collectively, these simulations indicate that Tld1 adopts significantly different conformational ensembles in the ER bilayer and LD environments, and that it interacts with TG molecules extensively in TG-rich LDs (**Fig 4E**). This provides a potential molecular explanation for Tld1 preferentially targeting to TG-rich LDs.

### Mutation of HR1 and HR2 residues alters LD targeting

Motivated by the MD simulations, we mutated key residues in the Tld1 HR1 and HR2 regions that were predicted to perturb LD targeting. Specifically, we mutated polar (K26E) and hydrophobic (F44D) residues on the HR1 amphipathic helix, as well as residues at the kink of the helix-kink-helix motif of HR2 (S76A, Q72A+S76A). As predicted, mutation of K26E or F44D resulted in total loss of detectable Tld1-GFP LD targeting in yeast (**Fig 5A**). A dim GFP signal was detected in the vacuole lumen, which may be from eventual protein degradation following the loss of LD targeting. Similarly, the Tld1-GFP Q72A+S76A mutant displayed significantly reduced LD targeting and a dim vacuole lumen signal (**Fig 5A**). However, the single S76A mutant demonstrated an intermediate effect, retaining some LD targeting. Overall, these mutations support the MD simulations and their proposed mechanism for how Tld1 targets to LDs.

**Figure 5.**
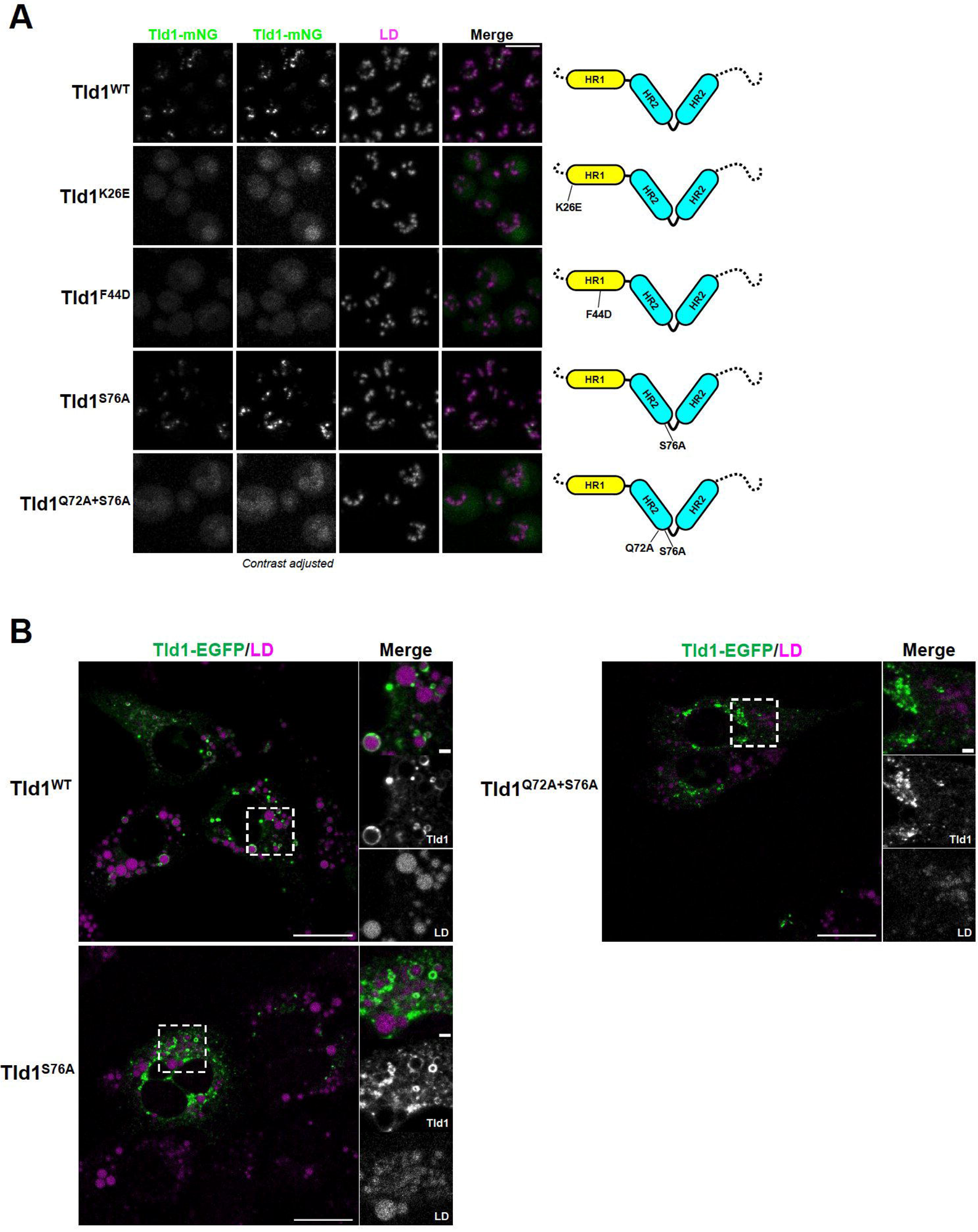
Point mutations in Tld1 HR regions disrupt LD-targeting in yeast and mammalian cells. **(A)** Imaging of yeast expressing full length Tld1 hydrophobic region (HR) point mutants tagged with mNG. LDs were stained with MDH (magenta). Dotted black lines on diagrams (far right) indicate N and C-termini of Tld1. Far left column represents non-contrast adjusted mNG channel. **(B)** Confocal imaging of U2-OS cells expressing either full length WT Tld1-EGFP or full length HR mutant Tld1-EGFP. LDs were stained with MDH (magenta). Images are midplane sections, with GFP and LD channels contrast adjusted. Yeast scale bar, 5µm. Mammalian scale bar, 20µm. Mammalian inset scale bar 2µm.

To test the impact of these mutations on the HR2 ensembles in simulations, we employed replica exchange umbrella sampling to calculate free energy profiles for kink formation in the ER bilayer and TG-rich LD (**Supplemental Fig 4A, 4B**). As anticipated, the wildtype profiles show a strong driving force for kink opening in the ER membrane and kink formation in the LD. Replacing both Gln72 and Ser76 with alanine (Q72A+S76A) not only reduces the kink-opening driving force in the ER membrane from ∼12 kcal/mol to ∼4 kcal/mol, but it also shifts the most stable conformation deeper into the membrane (∼1.5nm below the phosphate plane). Interestingly, the kink formation remains in the LD, verifying the kinked conformation is most stable when polar interactions for the remaining polar groups, including backbone interactions, are accessible in the LD monolayer. These free energy profiles verify that Gln72 and Ser76 play a dominant role in conformational change in HR2 upon LD targeting. Interestingly, the single S76A has an intermediate effect, reducing the kink-opening force in the ER membrane to ∼8 kcal/mol while retaining the kinked formation in the TG-rich LD. This would both decrease the kinetic barrier for LD targeting, consistent with the partially retained LD targeting for the S76A mutant. It could also increase stability in the ER membrane, though not significantly enough to inhibit experimental targeting like Q72A+S76A, consistent with the milder shift in the ER free energy profile (**Supplemental Fig 4A**).

Since the HR2 region appeared to mediate targeting of Tld1 to the ER network, we examined how the Q72A+S76A mutations impacted the localization of Tld1 to LDs and the ER network. To access this, we expressed Tld1-EGFP Q72A+S76A in HeLa cells treated with OA. Notably, whereas wildtype Tld1-EGFP concentrated on the surfaces of LDs, Tld1-EGFP Q72A+S76A localized primarily to large aggregates and was not generally detectable on LDs (**Fig 5B**). In contrast, Tld1-EGFP S76A showed some LD targeting but this appeared reduced compared to WT Tld1-EGFP. Collectively, this supports a model where both HR1 and HR2 contribute to Tld1’s LD association. We speculate that HR2 enables Tld1 to initially localize to the ER surface and engage the LD surface via HR1. Following LD localization, HR2 adopts a helix-kink-helix conformation that potentially reinforces LD localization together with HR1.

### Loss of Tld1 alters TG levels via enhanced TG lipolysis

What is the physiological function of Tld1? Because Tld1 LD targeting requires TG, and MD simulations indicate Tld1:TG interactions, we next determined whether manipulating Tld1 expression influenced cellular TG pools. We first examined steady-state TG and SE levels of WT and *tld1*Δ yeast. At LOG phase, *tld1*Δ yeast display a ∼20% steady-state reduction in TG compared to WT, while SE levels are unaffected (**Fig 6A**). We reasoned this TG reduction could be due to either enhanced lipolysis or decreased TG synthesis (or a combination of both).

**Figure 6.**
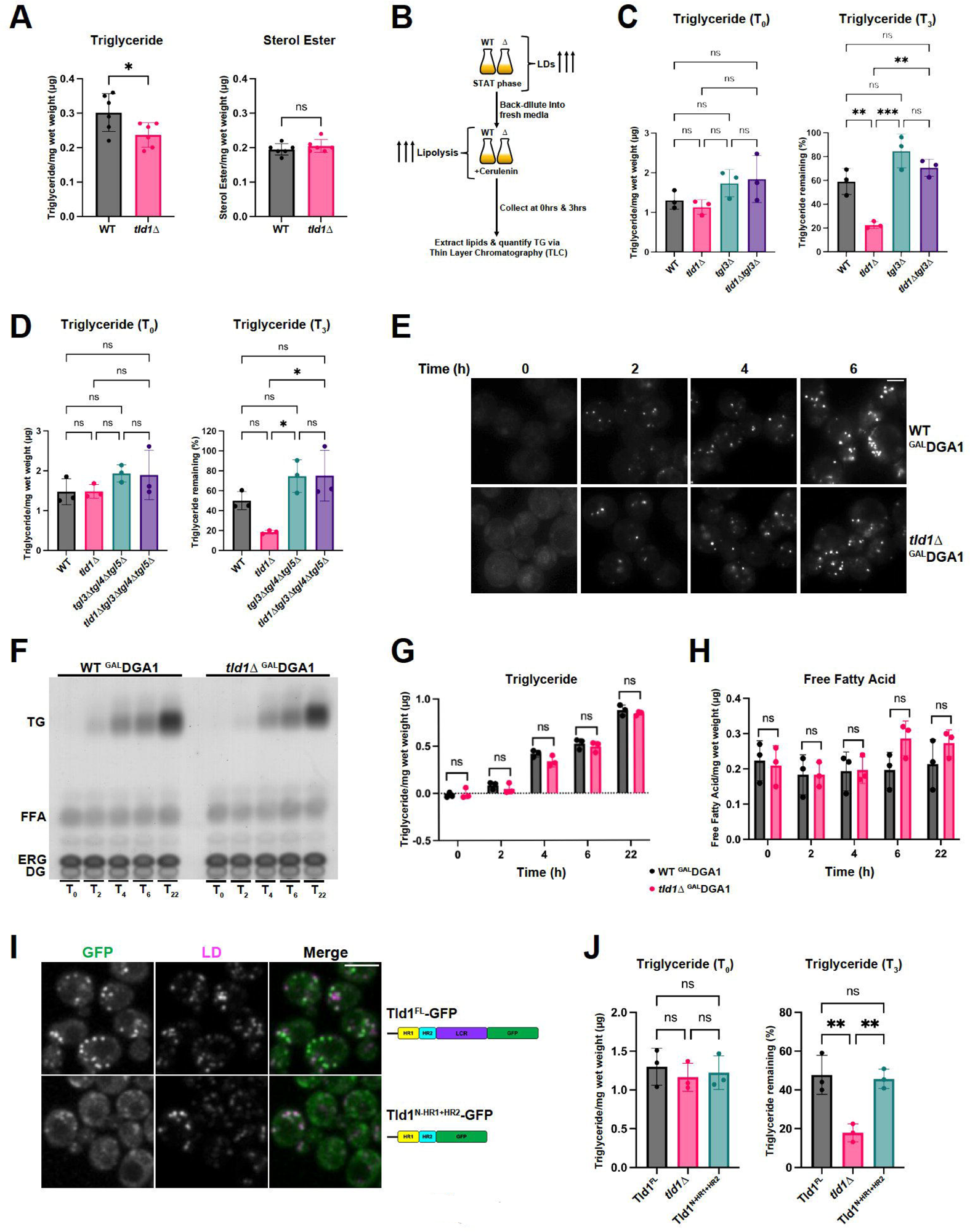
Tld1 deletion results in enhanced Tgl3 lipase-dependent TG lipolysis. **A)** Log phase, whole cell Triglyceride (TG, left graph) and Sterol Ester (SE, right graph) levels of wildtype (WT) and *tld1*Δ yeast, measured by TLC. Experiments conducted in triplicate. Statistical analysis is Unpaired t test with Welch’s correction. **(B)** Graphical schematic of cerulenin lipolysis assay for yeast. **(C, left panel)** TLC quantification of stationary phase, pre-lipolysis (T_0_) TG levels of WT, *tld1*Δ, *tgl3*Δ, and *tld1*Δ*tgl3*Δ. **(C, right panel)** Rate of lipolysis determined via TLC after treatment with 10µg/mL cerulenin (T_3_). Quantification represents percentage of starting TG that remained (pre-cerulenin TG levels set to 100% for each strain) after 3hrs of cerulenin-stimulated lipolysis. Experiments conducted in triplicate. Statistical analyses are ordinary one-way ANOVA. **(D, left panel)** TLC quantification of stationary phase, pre-lipolysis TG levels of WT, *tld1*Δ, *tgl3*Δ*tgl4*Δ *tgl5*Δ, and *tld1*Δ*tgl3*Δ*tgl4*Δ *tgl5*Δ. **(D, right panel)** Rate of lipolysis determined via TLC after addition of 10µg/mL cerulenin. Quantification represents percentage of starting TG that remained (pre-cerulenin TG levels set to 100% for each strain) after 3hrs of cerulenin-stimulated lipolysis. Experiments conducted in triplicate. Statistical analyses are ordinary one-way ANOVA. **(E)** Time-lapse imaging of galactose-induced LD formation in WT ^GAL^DGA1 and *tld1*Δ ^GAL^DGA1 yeast stained with MDH. Images are midplane sections. Scale bar 2µm. **(F)** Representative TLC plate of galactose-induced TG production in WT ^GAL^DGA1 and *tld1*Δ ^GAL^DGA1 yeast strains. FFA = Free Fatty Acids, ERG = Ergosterol, DG = Diacylglycerol. **(G)** TLC quantification of TG levels after galactose-induced TG production time-course in WT ^GAL^DGA1 and *tld1*Δ ^GAL^DGA1. Representative of three independent experiments. Statistical analyses are multiple unpaired t tests with Welch’s correction. **(H)** TLC quantification of FFA levels after galactose-induced TG production time-course in WT ^GAL^DGA1 and *tld1*Δ ^GAL^DGA1. Representative of three independent experiments. Statistical analyses are multiple unpaired t tests with Welch’s correction. **(I)** Imaging of endogenous WT full-length Tld1-GFP (Tld1^FL^-GFP) and truncated Tld1 with GFP inserted after HR2 (Tld1^N-HR1+HR2^-GFP), with MDH stained LDs. Scale bar, 5µm. **(J, left panel)** TLC quantification of stationary phase, pre-lipolysis TG levels of Tld1^FL^-GFP, *tld1*Δ, and Tld1^N-HR1+HR2^-GFP. **(J, right panel)** Rate of lipolysis determined via TLC after addition of 10µg/mL cerulenin. Quantification represents percentage of starting TG that remained (pre-cerulenin TG levels set to 100% for each strain) after 3hrs of cerulenin-stimulated lipolysis. Experiments conducted in triplicate. Statistical analyses are ordinary one-way ANOVA. *, P < 0.05; **, P < 0.01; ***, P < 0.001; ****, P < 0.0001.

To dissect this, we first tested whether TG lipolysis or TG biosynthesis were altered in *tld1*Δ yeast. Yeast contain three TG lipases: Tgl3, Tgl4, and Tgl5, of which Tgl3 performs the majority of the TG lipolysis (Athenstaedt and Daum, 2003, 2005). To determine whether TG lipolysis was altered in *tld1*Δ yeast, we treated WT, *tld1*Δ, *tgl3*Δ, and *tld1*Δ*tgl3*Δ yeast with cerulenin, which promotes TG lipolysis by blocking *de novo* fatty acid synthesis (making LDs the only source of fatty acids) (**Fig 6B**). We then measured yeast TG levels before (T_0_) and after 3hrs (T_3_) of cerulenin treatment. Importantly, WT and *tld1*Δ yeast contained similar TG levels at T_0_ because we allowed yeast to grow for 24hrs into STAT phase and accumulate TG (**Fig 6C**). After 3hrs of cerulenin, WT yeast had ∼60% of their TG stores remaining, whereas *tld1*Δ only had ∼20%, suggesting TG lipolysis was elevated in *tld1*Δ yeast (**Fig 6C**). In contrast to *tld1*Δ yeast*, tld1*Δ*tgl3*Δ yeast retained ∼70% of their TG, behaving similar to *tgl3*Δ yeast, suggesting the elevated TG loss in *tld1*Δ yeast required Tgl3. Since yeast also encode Tgl4 and Tgl5 TG lipases, we also performed similar experiments with WT, *tld1*Δ, *tgl3*Δ*tgl4*Δ*tgl5*Δ, and *tld1*Δ*tgl3*Δ*tgl4*Δ*tgl5*Δ yeast (**Fig 6D**). Of note, *tgl3*Δ*tgl4*Δ*tgl5*Δ yeast and *tld1*Δ*tgl3*Δ*tgl4*Δ*tgl5*Δ contained nearly identical TG levels following 3hrs of cerulenin-induced TG lipolysis. Collectively, this supports a model where *tld1*Δ yeast exhibit enhanced TG lipolysis that is suppressed by genetic depletion of Tgl lipases.

Next, we determined whether Tld1 loss alters TG biosynthesis. We utilized a yeast strain in which all of the acyltransferases that synthesize neutral lipids were deleted, with the exception of Dga1. In this strain, the *DGA1* gene was placed under a galactose inducible *GAL* promoter (*are1*Δ*are2*Δ*lro1*Δ^GAL^DGA1, referred here simply as ^GAL^DGA1) (Cartwright et al., 2015). As expected, in the absence of galactose, this yeast strain contains no neutral lipids and no LDs, and therefore staining yeast with MDH reveals no LD foci (**Fig 6E, time T=0**). In the presence of galactose in the growth media, yeast synthesize TG via Dga1 expression and activity. The *GAL* promoter also enables all strains in this background to express the same level of Dga1. We now determined whether Tld1 loss impacted TG accumulation in this system. We deleted *tld1*Δ in this strain (*tld1*Δ^GAL^DGA1) and compared it and the ^GAL^DGA1 strain’s abilities to produce LDs and TG. First, we imaged LDs via MDH stain, finding that the ^GAL^DGA1 and *tld1*Δ^GAL^DGA1 strains generated comparable numbers of LDs after galactose induction (**Fig 6E**). Next, we measured whole-cell TG levels in these strains following GAL induction of TG synthesis. We found no significant difference in TG between these strains over multiple time-points (**Fig 6F, 6G**). We also detected no significant changes in free fatty acids (FFA) for either strain, although there was a very slight increase in FFAs in the *tld1*Δ yeast after 6 hrs, potentially due to enhanced TG lipolysis (**Fig 6H**). Altogether, these results support a model where the absence of Tld1 does not impact stepwise TG synthesis, and supports a model where reduced TG in *tld1*Δ yeast is primarily due to enhanced lipolysis.

#### The Tld1 HR1 and HR2 regions are sufficient for Tld1 function

Since the Tld1 HR1 and HR2 regions appeared necessary for LD targeting, we next asked whether these regions were sufficient for Tld1 function. We generated yeast with chromosomally GFP-tagged full length Tld1 (Tld1^FL^-GFP), or truncated Tld1 lacking the low complexity region (LCR) (Tld1^N-HR1+HR2^-GFP). Both GFP-tagged strains localized to LDs, although Tld1^N-HR1+HR2^-GFP appeared slightly dimmer on LDs (**Fig 6I**). We then tested whether Tld1^N-HR1+HR2^-GFP yeast exhibited reduced TG levels compared to Tld1^FL^-GFP using the cerulenin-induced lipolysis assay. As expected, initial (T_0_) TG levels for Tld1^FL^-GFP, Tld1^N-HR1+HR2^-GFP, and *tld1*Δ yeast were not significantly different (**Fig 6J**). After 3hrs of cerulenin-stimulated lipolysis (T_3_), TG levels of Tld1^N-HR1+HR2^ yeast were similar for Tld1^FL^ yeast, and significantly higher than *tld1*Δ yeast (**Fig 6J**). This suggests that the C-terminal LCR is not necessary for Tld1 function.

### Tld1 over-expression results in TG accumulation and LD enlargement

Since Tld1 loss lowered cellular TG, we next determined how Tld1 over-expression would influence LD neutral lipids. We measured steady-state TG and SE levels of WT yeast expressing either an empty vector (EV) or over-expressed Tld1 (Tld1 OE) on a strong *GPD* promoter. Strikingly, we observed a more than ∼4-fold increase in TG in Tld1 OE yeast compared to EV controls (**Fig 7A**). Notably, there was no effect on SE levels, suggesting Tld1 OE selectively impacted TG pools (**Fig 7A)**. In line with this, we observed enlarged LDs in Tld1 OE when they were imaged by thin section transmission electron microscopy (TEM) (**Fig 7B**). Quantification of TEM micrographs confirmed significantly increased LD sizes and numbers of detected LDs per thin-section of Tld1 OE cells compared to WT (**Fig 7C, 7D**), suggesting Tld1 OE elevated TG stores that were stored in enlarged LDs. A portion of the LDs observed in Tld1 OE had similar area to those of EV LDs, which are likely explained by the varying expression levels of the Tld1 OE construct. Collectively, this indicates that Tld1 OE correlates with elevated TG levels and enlarged LDs.

**Figure 7.**
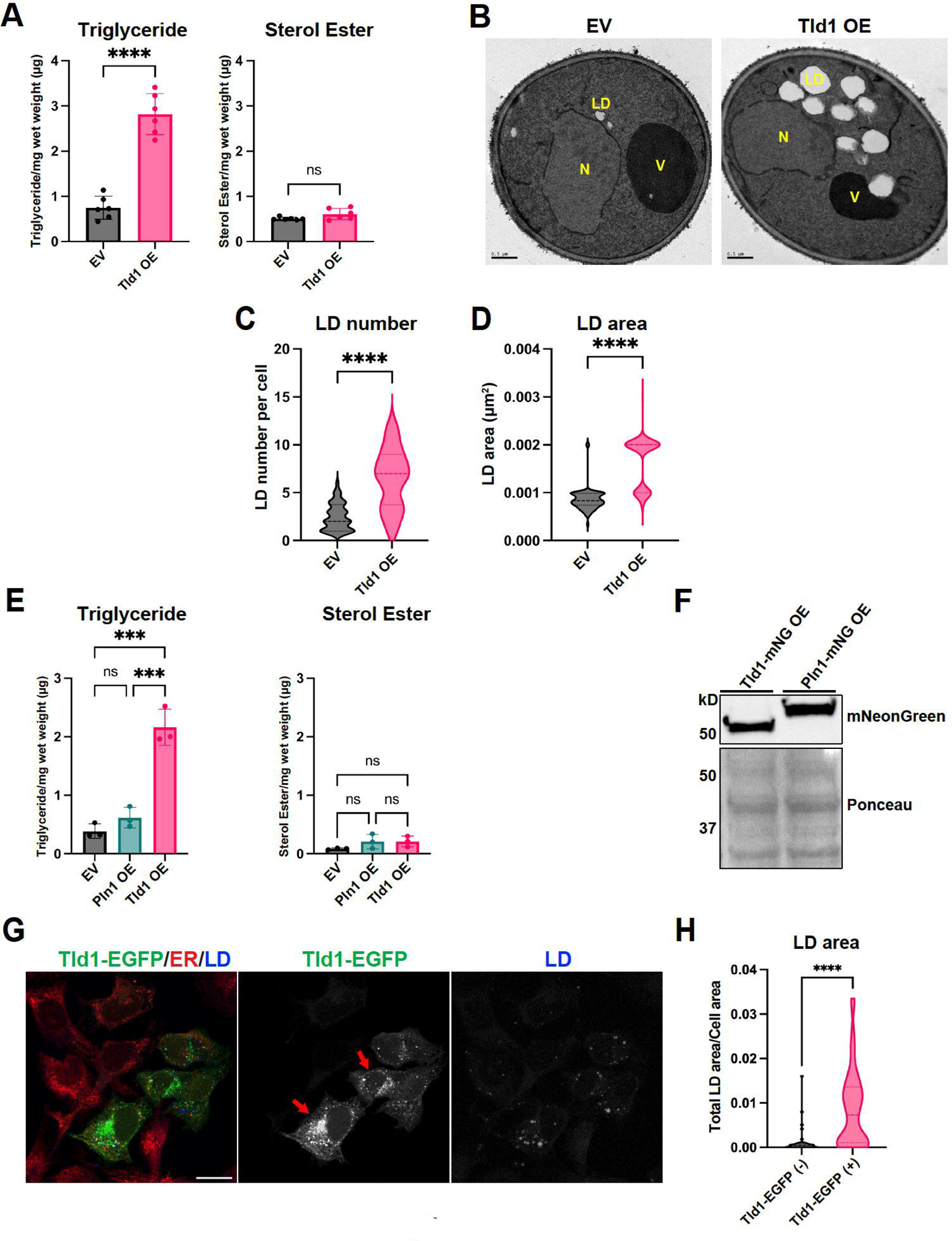
Overexpression of Tld1 significantly elevates TG levels, LD number and LD size. **(A)** Steady state, logarithmic phase Triglyceride (TG, left panel) and Sterol Ester (SE, right panel) levels in empty vector plus soluble GFP (EV) and Tld1-mNeonGreen overexpressing (Tld1 OE) yeast as quantified by TLC. Data were compiled from three independent experiments. **(B)** Thin-section TEM micrographs of logarithmic phase EV and Tld1 OE yeast. LD = Lipid Droplet, N = Nucleus, V = Vacuole. Scale bars, 0.5 µm**. (C)** LD number quantification from Fig 7B micrographs. n = 44 cells for EV and n = 18 cells for Tld1 OE. **(D)** LD area quantification from Fig 7B micrographs. n = 98 LDs for EV and n = 115 LDs for Tld1 OE. **(E)** Steady state TG (left panel) and SE (right panel) levels for EV, Tld1 OE, and Pln1 overexpressing (Pln1 OE) yeast. Experiments were performed in triplicate. **(F)** Protein expression of Tld1-mNeonGreen (Tld1-mNG OE) and Pln1-mNeonGreen (Pln1-mNG) overexpressing constructs used in Fig 7E. Membranes blotted with anti-mNeonGreen antibody and Ponceau S stain served as loading control for total protein. **(G)** Imaging of U2-OS cells overexpressing full length Tld1-EGFP. Cells were IF stained with α-HSP90B1 (ER, red), and LDs visualized with MDH (blue). Red arrows indicate cells with high expression of Tld1-EGFP. Scale bar, 20µm. **(H)** Quantification of average LD area per cell area from Tld1-EGFP negative cells (Tld1-EGFP (-)) and Tld1-EGFP positive cells (Tld1-EGFP (+)) from images in Fig 7G. n=39 cells for Tld1-EGFP (+) and n=48 cells for Tld1-EGFP (-). Statistics for Fig 7A, C, D, and H were unpaired t test with Welch’s correction. Statistics for Fig 7E was ordinary one-way ANOVA. *, P < 0.05; ***, P < 0.001; ****, P < 0.0001.

A possible explanation for the TG accumulation in Tld1 OE yeast is simply from over-expressing a LD surface protein, which could potentially crowd away other LD-resident proteins and indirectly perturb TG homeostasis (Kory et al., 2015). To test this possibility, we measured steady-state neutral lipids of yeast over-expressing Pln1 (also known as Pet10), a well characterized yeast perilipin-like protein (Gao et al., 2017), and compared these to EV and Tld1 OE expressing yeast (**Fig 7E**). Strikingly, Pln1 OE did not alter TG levels, which closely mirrored the EV control, and did not phenocopy the TG accumulation observed with Tld1 OE (**Fig 7E**). Notably, neither of the constructs altered SE pools. This indicated that the TG accumulation caused by Tld1 OE was likely not an artifact of simply overexpressing any LD protein, and supported a model where Tld1 OE specifically influenced LD TG pools. In further support, Western blot analysis of Tld1 OE and Pln1 OE expression levels revealed very similar expression levels of both proteins, suggesting they were expressing at similar high levels (**Fig 7F**). Collectively, this supports a model where Tld1 influences LD TG pools, and that its over-expression is sufficient to promote TG accumulation.

We also determined whether Tld1 over-expression in human cells affected LD sizes. We over-expressed Tld1-EGFP (Tld1-EGFP OE) in human U2-OS cells not treated with OA, since these cells typically have small but detectable LDs. Notably, Tld1-EGFP OE increased total LD size per cell area compared to control cells (**Fig 7G, 7H, red arrows are Tld1-EGFP positive cells**). This further suggests that, similar to yeast, Tld1 OE is sufficient to increase LD size.

### Tld1 loss or over-expression does not impact Tgl lipase abundance nor LD targeting

How does Tld1 influence LD TG pools? One possibility is that Tld1 loss or over-expression may alter the total abundance or LD localization of TG lipases. To test this, we performed fluorescence imaging of GFP-tagged TG lipases Tgl3, Tgl4, and Tgl5 in WT and *tld1*Δ yeast (**Fig 8A**). Imaging revealed there were no obvious changes in Tgl lipase LD targeting in the absence of Tld1, suggesting Tgl LD targeting was intact in *tld1*Δ yeast. We also examined Tgl protein levels by Western blotting. Steady-state protein abundances of Tgl3, Tgl4, and Tgl5 were unaffected by Tld1 loss, indicating the enhanced lipolysis observed in *tld1*Δ yeast was not simply due to increased total lipase abundances (**Fig 8B**).

**Figure 8.**
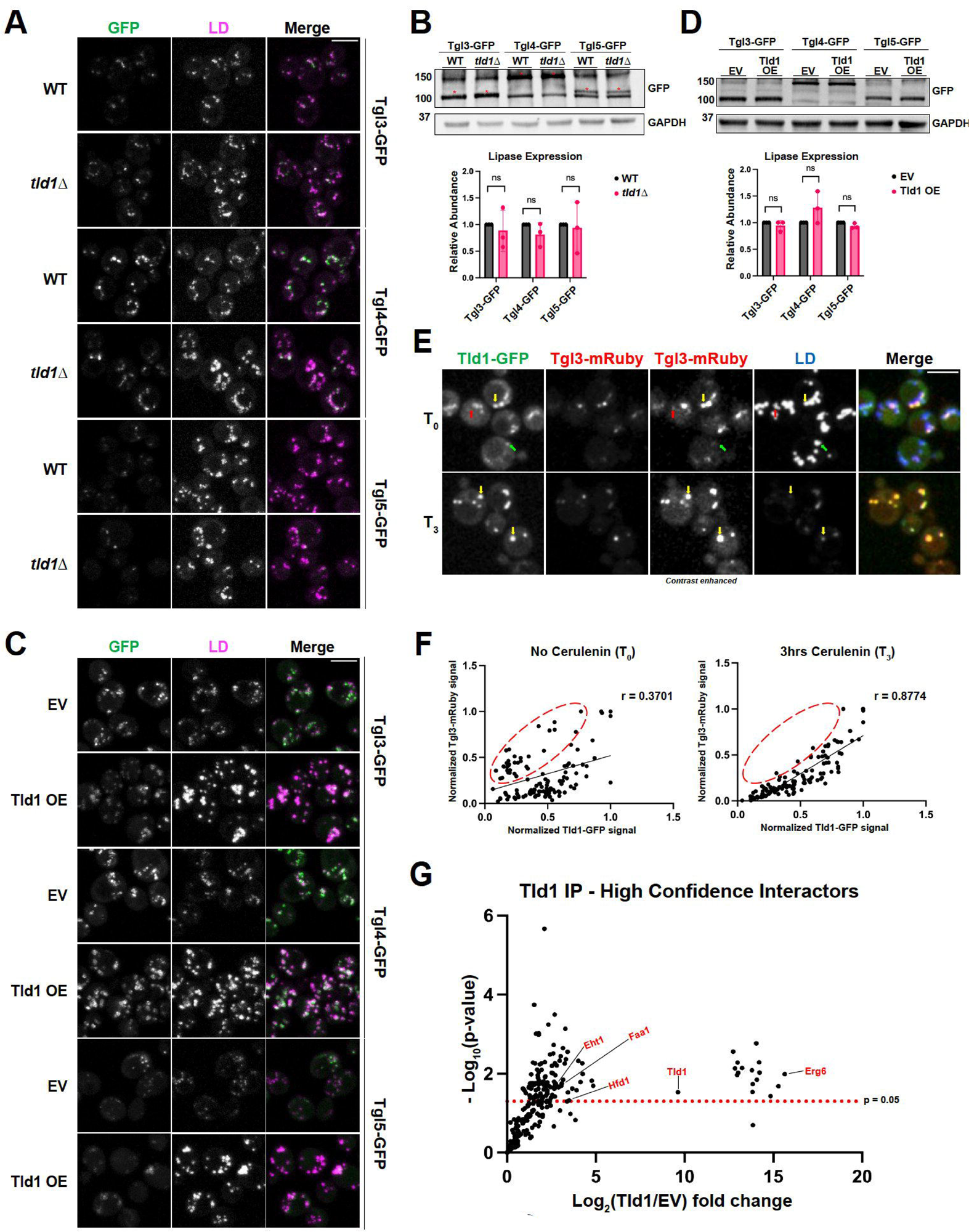
Tld1 does not alter TG lipase LD-targeting or protein abundance, but modulates lipolysis on the LD. **(A, C)** Fluorescence imaging of GFP-tagged TG lipases in either WT and *tld1*Δ (A) or EV and untagged Tld1 OE yeast (C). LDs were stained with MDH. **(B, D)** Protein expression levels of GFP-tagged TG lipases in WT and *tld1*Δ (B) and EV and Tld1 OE (D). Red asterisks indicated GFP-tagged lipases. Data is normalized to WT or EV, respectively, and represents three independent experiments. **(E)** Fluorescence imaging of Tld1-GFP and Tgl3-mRuby dual-tagged yeast, with MDH stained LDs before (T_0_) and 3hrs after cerulenin-stimulated lipolysis (T_3_). Green arrows indicate Tld1-enriched LDs, red arrows indicate Tgl3-enriched LDs, and yellow arrows indicate LDs targeted with both Tld1 and Tgl3. Second from left column represents non-contrast adjusted images for Tgl3-mRuby. **(F)** Scatterplot of Tld1-GFP fluorescence signal intensity versus Tgl3-mRuby signal intensity for random LDs before (T_0_) and after 3hrs cerulenin treatment (T_3_) with Pearson’s correlation coefficient (r) displayed. Data corresponding to images in Fig 8E. Red circles indicate Tgl3-enriched/Tld1-deenriched LDs. n = 120 LDs for each condition, quantified from 87 cells for T_0_ and 105 cells for T_3_. **(G)** Volcano plot showing negative Log_10_ P value (-Log_10_) and Log_2_ abundance changes for Tld1 IP interactors versus EV control, obtained via mass spec analysis. Red text at select data points indicates LD proteins found to directly interact with Tld1. Red dotted line indicates significance cut-off for protein hits. Data were collected from three independent experiments. Statistical analyses are multiple unpaired t tests. Scale bars, 5µm. ns, ≥ 0.05.

We also determined whether Tgl1 OE affected Tgl abundances or localizations. As expected, yeast over-expressing Tld1 displayed larger and more numerous LDs, but this did not alter the LD localization of any of the GFP-tagged Tgl proteins, suggesting Tld1 OE does not inhibit their LD targeting (**Fig 8C**). We did note a slight change in the fluorescence pattern of Tgl lipases in Tld1 OE cells, but reasoned this may be due to increased LD sizes, which would distribute Tgl lipases across a larger LD surface area. In line with this, Western blotting revealed no significant changes in the abundances of Tgl lipases in Tld1 OE yeast compared to WT, indicating that the TG accumulation was not due to decreased lipase expression (**Fig 8D**). Collectively, this indicates that changes in steady-state TG levels in *tld1*Δ or Tld1 OE are not due to changes in the abundances of TG lipases or their LD targeting.

To determine whether Tld1 may physically interact with Tgl lipases on the LD surface, we also conducted co-immunoprecipitation (co-IP) experiments where we over-expressed either mNG (EV-mNG) alone or Tld1-mNG in yeast, immunoprecipitated with anti:mNG affinity resin, and examined the co-IP fractions by LC-MS/MS proteomics. Notably, numerous canonical LD proteins were significantly enriched in the Tld1-mNG co-IP fraction, including Erg6, Hfd1, Faa1, and Eht1 (**Fig 8G**). However, we did not detect any peptides from Tgl3, Tgl4, nor Tgl5 in this experiment. Similarly, we conducted co-IPs followed by Western blotting of Tld1-mNG yeast also expressing 3xHA-tagged Tgl3. While we detected Tgl3-3xHA in the input, we did not detect it in the Tld1-mNG co-IP fraction (**Supplemental Figure 5A**). Collectively, this suggests that Tld1 and Tgl lipases do not form strong, stable physical interactions. However, we cannot rule that Tld1 and Tgl lipases form lower affinity interactions that may fine-tune or influence Tgl lipase activity on the LD surface.

#### Tld1 and Tgl3 independently target to LD subsets

Since Tld1 manipulation did not appear to influence Tgl lipase abundance or localization, we hypothesized that Tld1 may demarcate a subset of TG-positive LDs, and interact with LDs independently of Tgl lipases. In this model, Tld1 may function as a negative regulator of TG lipolysis on these Tld1-positive LDs, which would exhibit resistance to lipolysis compared to Tld1-negative LDs.

To test this model, we directly compared Tld1 and Tgl3 LD localizations in yeast co-expressing chromosomally tagged Tld1-GFP and Tgl3-mRuby. Prior to cerulenin treatment (T_0_), we observed LDs with detectable levels of both Tld1-GFP and Tgl3-mRuby (**Fig 8E, yellow arrows**), as well as LDs exhibiting only detectable Tld1-GFP (**Fig 8E, green arrows**) or Tgl3-mRuby alone (**Fig 8E, red arrows**). This indicates that Tld1 and Tgl3 can localize to the same LD, but also decorate separate LD subsets within a cell, suggesting they target to LDs independently of one another.

Next, we imaged this dual-labeled strain following 3hrs (T_3_) of cerulenin treatment to induce lipolysis (**Fig 8E**). We then quantified individual LD signals for Tld1-GFP and Tgl3-mRuby signal, and generated signal correlation graphs (**Fig 8F**). At T_0_ there is a heterogenous mix of Tld1-GFP and Tgl3-mRuby signals on LDs, with some LDs displaying abundant Tgl3-mRuby signal but low Tld1-GFP signal (Tgl3>Tld1, upper-left region of chart, red circle), LDs with significant levels of both Tld1-GFP and Tgl3-mRuby signals (Tgl3∼Tld1, center to upper-right region of chart), and LDs with high Tld1-GFP but low Tgl3-mRuby signal (Tld1>Tgl3, lower right region of chart). The signals were poorly correlated with a Pearson correlation of r=0.3701, supporting the model where Tld1 and Tgl3 target to LDs independently. However, following 3hrs of cerulenin treatment, the Tld1-GFP/Tgl3-mRuby LD signal distribution changed. LDs now displayed a more linear positive correlation pattern, with Tld1-GFP signal better correlating with Tgl3-mRuby signal (i.e., Tgl3∼Tld1). This is reflected with a significantly higher Pearson correlation of r=0.8774 (**Fig 8F**). Notably, yeast with low Tld1-GFP signal and high Tgl3-mRuby were generally absent following cerulenin treatment, supporting a model where they were lost during lipolysis (**Fig 8F, red circle**). Collectively, this supports a model where Tld1-positive LDs may be more resistant to complete mobilization during lipolysis.

### Tld1 loss alters LD accumulation in yeast stationary phase

As yeast transition into STAT phase, they enter slow growth and shunt excess lipids into TG for long-term storage. Since Tld1 loss elevated TG lipolysis, we queried whether *tld1*Δ yeast would display differences in LD abundances as they transitioned into STAT phase. We quantified the number of LDs per yeast cell for WT and *tld1*Δ yeast initially cultured in 2% glucose media and allowed to grow continually in this media for six days (defined as gradual glucose restriction, GGR). At the start of the experiment (T=0 days), when cells were in early STAT phase, *tld1*Δ yeast exhibited more MDH-stained LDs compared to WT (**Fig 9A, 9B**). However, following six days of GGR, *tld1*Δ yeast displayed significantly fewer LDs per cell than WT yeast (**Fig 9A, 9B**). This supports a model where Tld1 depletion causes elevated TG lipolysis, which over time would gradually deplete LD stores in yeast subsisting in low-nutrient conditions. Collectively, we propose a model in which Tld1 labels a subset of TG-containing LDs and marks them for preservation from lipolysis, which could in principle be utilized as a lipid source in stationary phase subsistence (**Fig 9C**).

**Figure 9.**
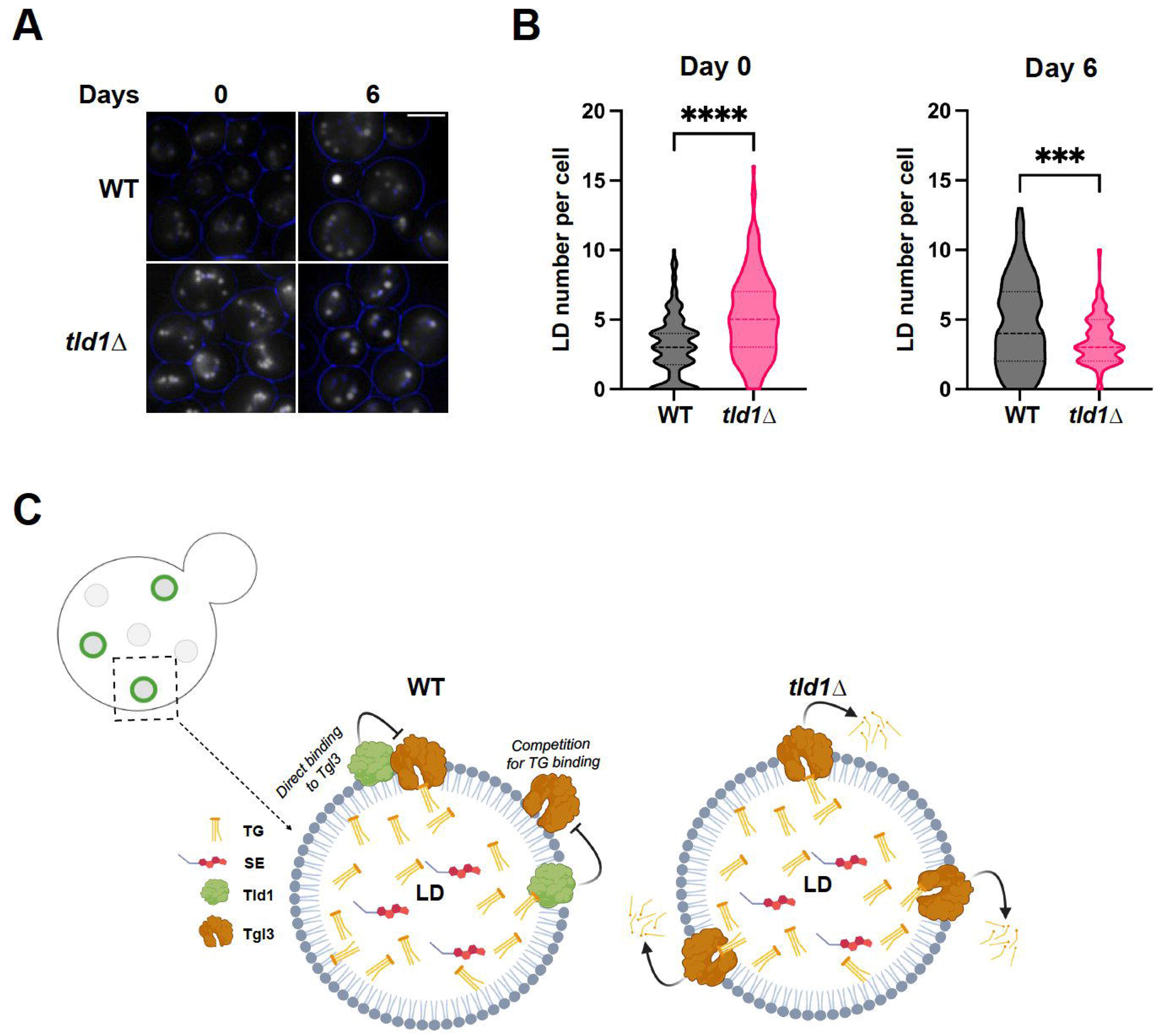
Tld1 influences LD maintenance during late starvation conditions. **(A)** Imaging of MDH-stained WT and *tld1*Δ yeast LDs, before (Day 0) and after (Day 6) exposure to late stationary phase. Blue circles indicate cell borders. Images are midplane sections. **(B)** Quantification of LD number per cell at Day 0 and Day 6 of exposure to starvation, from images in Fig 9A. n = 150 cells for both WT and *tld1*Δ, each. **(C)** Cartoon working model of Tld1 negative regulation of Tgl3-dependent TG lipolysis. Upper left depicts a yeast cell with Tld1 negative LDs (gray circles) and Tld1 positive LDs (green rings on gray circles). Black box indicates a representative Tld1-positive LD and arrow points to close up of working model. This may occur through competition for TG substrate binding or a weak protein-protein interaction (WT LD, bottom left). In the absence of Tld1, TG is more accessible to Tgl3 lipase (*tld1*Δ LD, bottom right). Statistics are unpaired t test with Welch’s correction. Scale bars, 5µm. **, P < 0.01; ****, P < 0.0001.

### Limitations of study

This study shows that Tld1-GFP localizes to subsets of yeast LDs, and requires TG synthesis for LD localization. This LD preference occurs independent of other yeast proteins, as Tld1 localizes to TG-rich LDs when ectopically expressed in human cells. Tld1 LD targeting requires HR1 and HR2, which work cooperatively for LD localization. Functionally we find that Tld1 influences TG pools in yeast. Tld1 loss correlates with enhanced TG lipolysis, whereas over-expressing Tld1 is sufficient to drive TG accumulation and LD enlargement. Collectively, this supports a model where Tld1 regulates Tgl lipase-dependent TG lipolysis. At least two mechanisms could explain how Tld1 inhibits lipolysis: 1) Tld1 may limit TG accessibility by altering LD surface properties and thereby the nature of lipase activity, or 2) Tld1 may bind directly to lipases or to intermediate molecules in a macromolecular complex to inhibit lipase enzymatic activity. Future studies will dissect the mechanism of Tld1-mediated TG interactions and lipolysis regulation.

## Discussion

LDs can be classified into distinct subpopulations, and these LD subsets are differentiated by unique proteomes, morphologies, or spatial distributions (Thiam and Beller, 2017; Eisenberg-Bord et al., 2018; Teixeira et al., 2018; Schott et al., 2019; Ugrankar et al., 2019). Two key questions are how proteins target to LD subsets, and how distinct proteomes confer specific functions to them. Here, we demonstrate that Tld1 is a yeast protein that enriches on a subpopulation of TG-containing LDs and acts as a negative regulator of TG lipolysis. We find that Tld1 LD targeting requires TG, and MD simulations reveal that Tld1’s two hydrophobic regions adopt specific conformations on TG-rich LDs that engage in TG interactions. Loss of Tld1 reduces steady state TG levels at LOG growth phase but does not alter SE pools. We find this is due to enhanced lipolysis. Tld1 over-expression promotes TG accumulation and LD enlargement, a phenotype not replicated by overexpressing another LD coat protein Pln1/Pet10. We also find that yeast lacking Tld1 display altered LD mobilization after multiple days in STAT phase. This may indicate that in the absence of Tld1, lipolysis is dysregulated and yeast struggle to maintain LD pools after long-term starvation. This supports a model where Tld1 helps yeast maintain a pool of TG-rich LDs to survive extended nutrient deprivation.

A pervasive question in LD biology is how proteins target to LDs. One factor that clearly influences both LD protein targeting and stability is the presence of neutral lipids in LDs (Grillitsch et al., 2011; Schmidt et al., 2013; Klein et al., 2016; Gao et al., 2017; Prévost et al., 2018; Chorlay and Thiam, 2020; Rogers et al., 2022). Our Tld1 structure-function analysis indicate that both HR1 and HR2 contribute to organelle targeting, and mutations to either HR1 or HR2 reduce LD targeting. In line with this, MD simulations indicate that HR1 and HR2 undergo significant conformation changes in response to different lipid environments. On LDs, HR2 adopts a more compact helix-kink-helix conformation and interacts with TG, in contrast to a more “splayed open” conformation in the ER bilayer. HR1 interacts extensively with TG and PLs on the LD surface, but disengages from the SE-rich LD surface, likely weakening Tld1 LD affinity. This indicates both HR1 and HR2 enable Tld1 to anchor on LDs, but HR1 may act as a LD “lipid composition sensor”, preferentially engaging TG-rich LDs and potentially explaining why Tld1 is detected on only LD subsets. In the absence of TG, Tld1 adopts a less favorable conformation and may be targeted for degradation, although further studies are needed to dissect this.

How does Tld1 regulate TG lipolysis? We show that Tld1 loss or over-expression does not alter Tgl lipase LD targeting nor protein abundance. We fail to detect strong interactions between Tgls and Tld1 by co-IPs and mass spectrometry analysis, indicating that Tld1 does not strongly interact with Tgls. While this indicates that Tld1 does not tightly interact with lipases, we cannot rule out that it influences lipase activity via weak allosteric interactions. An alternative hypothesis is that Tld1 alters the LD surface properties in a manner that alters TG access to lipases. Further work is needed to fully dissect how Tld1 influences lipolysis.

Previous visual screens have identified a number of yeast proteins that target LD subsets. The most studied examples are the isoforms Ldo16 and Ldo45, which demarcate a specific pool of LDs near the yeast nucleus-vacuole junction (NVJ). These NVJ-associated LDs also contain other LD proteins like Pdr16 (Eisenberg-Bord et al., 2018; Teixeira et al., 2018; Ren et al., 2014). In their investigation, Eisenberg-Bord et al also identified Tld1 (called Bsc2 at that time) as a marker of the Pdr16-enriched LD subset (Eisenberg-Bord et al., 2018). Our study now characterizes Tld1 as a negative regulator of TG lipolysis, suggesting that NVJ-associated LDs may be protected from lipolysis so they can serve as substrates for lipophagy near the NVJ during long-term starvation. In the future, we hope to further reveal the function of this Tld1-positive LD subset in yeast physiology and metabolic adaptation.

## Acknowledgements

The authors would like to thank members of the Henne and Swanson labs for conceptual help on this manuscript. We also thank Jonathan Friedman, Joel Goodman, and Daniela Nicastro for conceptual assistance. We would also like to thank Joel Goodman for providing W303 yeast strains for this study. Finally, we would like to thank the UT Southwestern proteomics, electron microscopy, and live cell imaging core facilities for their aid with data collection and analysis. W.M.H, N.O.S. and E.G.R. are supported by grants from the Welch Foundation (I-1873), the NIH NIGMS (GM119768), NIDDK (DK126887), the Ara Parseghian Medical Research Fund, and the UT Southwestern Endowed Scholars Program. J.M.J.S. and R.J.B are supported by NIH NIGMS (R35GM143117) and are grateful for computational support from ANTON 2 at the Pittsburgh Supercomputing Center (MCB210016P), EXPANSE at the San Diego Supercomputing Center through the ACCESS program (allocation MCB200018) supported by NSF (grants #2138259, #2138286, #2138307, #2137603, and #2138296), as well as the Center for High Performance Computing at the University of Utah.

## Methods

### Yeast growth conditions

The WT parental strain used for all experiments and cloning in this study was BY4741 (MATa *his3*Δ1 *leu2*Δ0 *met15*Δ0 *ura3*Δ0). W303-1A (MATa leu2-3, 112 trp1-1 can1-100 ura3-1 ade2-1 his3-11,14) yeast strain was used as the parental strain for imaging of yeast with different neutral lipid-containing backgrounds (Fig 1E,F) and for galactose induced TG synthesis experiments (Fig 6E-H). Synthetic-complete (SC) growth media was used for culturing yeast cells in all experiments, except for experiments where uracil was excluded to retain pBP73G and pRS316 plasmids, or leucine excluded to retain pRS305 plasmid. For all experiments (unless noted below), a colony of yeast was inoculated from a YPD (yeast extract peptone dextrose) plate into SCD (SC dextrose/glucose) media and allowed to grow for ∼ 24hr in a 30°C incubator with shaking at 210 rpm. These cultures were diluted to OD_600_ = 0.001 in SCD media containing 2% glucose (wt/vol), grown overnight in a 30°C incubator shaking at 210 rpm, and collected at mid-log phase (∼OD_600_ = 0.6) the next day. For cerulenin experiments, yeast were cut back to OD_600_ = 0.1 in SCD media from an overnight culture and grown for 24hr, 30°C, 210 RPM. 50 OD_600_ units were collected from the 24hr culture as pre-lipolysis sample (Time 0hrs, “T_0_”). The remainder of the 24hr culture was cut back to OD_600_ = 0.5 in fresh SCD media containing 10µg/mL Cerulenin (Cat# C2389; Sigma-Aldrich) final concentration and allowed to incubate for 3hrs before harvesting 50 OD_600_ units as a post-lipolysis sample (Time 3hrs, “T_3_”). Aliquots were then washed in MiliQ water, pelleted, and then processed for lipid extraction and TLC. Culturing of yeast for cerulenin imaging experiments (Fig 8E) was done as detailed above, except a small aliquot was removed from 24hr culture as T_0_ and ∼25 OD_600_ was removed at T_3_ from SCD plus cerulenin cultures. All samples were concentrated down to 1mL in their respective media, and LDs stained for 5 min with MDH (SM1000a; Abcepta) at a final concentration of 0.1 mM before imaging. For induction of TG synthesis, ^GAL^DGA1 yeast strains were first cultured in 0.2% dextrose SCD media, overnight. Cells were then pelleted, washed in MiliQ water, and resuspended in 2% raffinose SCR media (2% raffinose substituted for dextrose in SCD) at OD_600_ = 0.5 and cultured for 24hrs. Following 24hr incubation, 50 OD_600_ units were removed as Time 0 (“T_0_”) sample for lipid extraction and TLC. The remainder of the yeast were pelleted, washed in MiliQ water, then cut back to OD_600_ = 2 in SCG media (2% galactose substituted for dextrose in SCD), and incubated for 22hrs. 50 OD_600_ unit aliquots were removed at 2, 4, 6, and 22hrs incubation, washed in Mili-Q water, then pelleted and processed for lipid extraction and TLC. For imaging of induced LDs, ^GAL^DGA1 yeast strains were cultured same as above, except 1mL aliquots were taken from SCG cultures at indicated time points and incubated for 5 min with MDH at a final concentration of 0.1 mM to visualize LDs. For gradual glucose restriction LD imaging experiments, yeast were cultured from a plate overnight in SCD media. 1mL of overnight culture was taken for Day 0, stained with MDH and LDs were imaged. The remainder of the culture was cut back to OD_600_ = 0.1 in fresh SCD media and incubated for 6 days, with 1mL aliquots taken each day and LDs imaged after staining with MDH. For immunoprecipitation and proteomics, cells were cultured from plates into SCD-URA (without uracil) overnight. Then, yeast were cut back to OD_600_ = 0.1 into same drop-out media, then incubated for 24hrs (under general growth conditions described above) until they reached stationary phase. After 24hrs, 250 OD_600_ units were collected for each sample, pelleted at 4000 RPM for 5 min, washed in Mili-Q water then pelleted again. Final yeast pellets were then subjected for protein extraction and immunoprecipitation.

### Molecular Dynamics Simulations

#### Structure prediction

TOPCONS (Tsirigos et al., 2015) and TmAlphaFold (Dobson et al., 2023) were used to predict the membrane-embedded regions of Tld1. The protein structure prediction tools RoseTTAFold (Baek et al., 2021) and AlphaFold2 (Jumper et al., 2021) were then used to model the structure of Tld1^N-HR1+HR2^ (amino acids 1-100). The resulting output poses from both resources agreed on the placement and alignment of all helices within the protein. This included HR1 in a single amphipathic helical structure and HR2 in a helix-kink-helix structure. The final structure was taken from RoseTTAFold, using no pairing or templates. Notably, the 5 top-scoring structures from RoseTTAFold had quite similar alignment. The output for TOPCONS transmembrane topology and the selected final structure are in Supplemental Figures 2M and 2C, respectively.

#### Simulations

The CHARMM36 force field (Huang et al., 2017) was used in all simulations. The bilayer system was created in the CHARMM-GUI membrane builder (Jo et al., 2008) with a ratio of 88:37:10 ratio of 3-palmitoyl-2-oleoyl-D-glycero-1-phosphatidylcholine (POPC), 2,3-dioleoyl-D-glycero-1-phosphatidylethanolamine (DOPE), and phosphatidylinositol (SAPI), respectively. A total of 585 PLs per leaflet (1170 PLs total) were used. The LD systems had the same 88:37:10 membrane composition ratios for their respective monolayer leaflets, for a total 135 PLs per leaflet (270 PLs total) and included an 8 nm thick neutral lipid core composed of a 90:10 CHYO:TG ratio for the SE-rich LD and a pure-TG core for the TG-rich LD (483 and 428 NLs respectively). These LD structures were taken from the last frame of 8 µs long simulations conducted in our previous work, which importantly had already obtained the properly equilibrated distributions (Braun and Swanson, 2022). Previous work has implemented the reduction of the partial charges of the glycerol moiety for TG in order to account for its bulk behavior such as interfacial surface tension (Kim et al., 2022; Campomanes et al., 2021). However, these models fail to accurately describe the area-per-lipid and packing defects of the PL monolayer surrounding the TG-core when simulated in an LD system (Kim et al., 2022). The TG forcefield parameters were obtained from our previous work (Kim and Swanson, 2020), in order accurately retain these properties that are important for protein-LD interactions. The membrane systems were embedded in 5nm of water and 0.15M NaCl on top and bottom to account for proper hydration and physiological conditions. To insert the Tld1 structure into the membrane systems, in-house MDAnalysis (Gowers et al., 2016) scripting was used, placing HR2 into the bilayer and LD monolayers and HR1 0.5 nm above the membrane. Overlapping PLs and neutral lipids were removed and the systems (9 PLs and 3 NLs for pure-TG LD, 9 PLs and 2 NLs for 90:10 LD, and 10 PLs from bilayer system). The systems were minimized for 5000 steps before being re-equilibrated for 10 ns using NVT conditions and 100 ns using NPT conditions. For the bilayer and TG-LD systems, long-timescale simulations lasting 4.5 µs were conducted using the Anton2 supercomputer provided by Pittsburg Supercomputing Center (Shaw et al., 2014), while the 90:10 CHYO:TG system was run for 1 μs on the EXPANSE supercomputer provided by San Diego Supercomputing Center (Strande et al., 2021). The Anton2 simulations were conducted using a 2.4 fs timestep, with the temperature set to 310 K, using Nose-Hoover thermostat (Nosé, 1984), and the MTK barostat (Martyna, Tobias, and Klein, 1994). Hydrogen bonds constrained with M-SHAKE (Kräutler et al., 2001), and the Gaussian-split Ewald was used to calculate long-range electrostatics (Shan et al., 2005). The additional parameters and intervals were used as suggested by the Anton2 website (https://wiki.psc.edu/twiki/view/Anton). EXPANSE simulations used a 2 fs timestep. The temperatures were set to 310 K, using Nose-Hoover thermostat (Nosé, 1984), (Hoover, 1985) and a temperature coupling time constant of 1 ps. The particle mesh Ewald (PME) algorithm (Essmann et al., 1995) was used to calculate the long-range electrostatic interactions with a cutoff of 1.0 nm. Lennard–Jones pair interactions were cutoff at 12 Å with a force-switching function between 8 and 12 Å, and pressure was maintained semi-isotropically using the Parrinello-Rahman barostat (Parrinello and Rahman, 1981). The pressure was set to 1.0 bar, with a compressibility of, 4 x 10^-S^ bar^-1^, and a coupling constant of 5.0 ps. The hydrogen bonds were constrained with the LINCS algorithm (Hess, 2008). We calculated the coordination numbers, RDFs, and protein positions using MDAnalysis, and in-house Python scripting, and Gromacs tools (Abraham et al., 2015), and the images were rendered using Visual Molecular Dynamics (VMD) (Humphrey et al., 1996).

##### Umbrella Sampling

PMFs for the positioning of proteins in the LD monolayer and ER bilayer were calculated using replica-exchange (Sugita and Okamoto, 1999) umbrella sampling simulations. The PMFs were calculated as a function of the z-coordinate center of mass (COM) of the 4 residues in the kink region of the HR2 sequence (residues 71-76). The windows were set up at an interval of 0.15 nm, spanning 0.7-2.5 nm below the PL-phosphate plane of the respective leaflets. The initial structures were initiated with steered MD simulations that were pulled at a rate of 0.001 nm/ps, with a force constant of 1000 kJ/mol/nm^2^. A harmonic umbrella potential of 1000 kJ/mol/nm^2^ was used for the umbrella sampling as well. The exchanged was attempted every 100 steps. After 70 ns of equilibration, the PMF was calculated using WHAM (Kumar et al., 1992) of the average of the last three-10 ns trajectories. The error bars were obtained from the standard deviation of these last 3 blocks. Umbrella sampling simulation were run using Gromacs version 2022.3 (Abraham et al., 2015).

#### Metadynamics

Potentials of mean force (PMFs) for single amino acids permeating through a bilayer were conducted using Well-Tempered Metadynamics (Barducci et al., 2008) biasing the *z*-component connecting the center of mass of the membrane and the center of mass of the amino acid. The bilayers used for the metadynamics simulations were created from same initial systems described above. The system was hydrated 5 nm of water surrounding each side with 0.15 M NaCl, and the respective amino acid was placed 2 nm above the membrane surface. The amino acids included in our simulations were Phe, Gln, Leu, Ser. The amino acids were neutralized by patching with the NH2 (CT2) group at the C-terminus, and an acetyl (ACE) at the N-terminus. Four replicas of each amino acid system were run for 500 ns each. The final PMF was obtained by averaging the PMFs obtained from the four simulations. The Gaussian function was deposited every 2 ps with a height of 0.05 kJ/mol and the bias factor was set to 15. Simulations were conducted in the canonical ensemble (NVT) at a temperature of 310K, using the Gromacs version 2019.4 (Abraham et al., 2015) patched with PLUMED version 2.5.3 (Tribello et al., 2014).

### Lipid extraction and TLC

For lipid extraction, 50 OD_600_ units of cells were collected for each sample, and pellet wet weights were normalized and recorded prior to extraction. Lipid extraction was performed using a modified Folch method (Folch et al., 1957). Briefly, cell pellets were resuspended in Milli-Q water with 0.5-mm glass beads (Cat # G8772-500G; Milipore Sigma) and lysed by three 1-min cycles on a MiniBeadBeater. Chloroform and methanol were added to the lysate to achieve a 2:1:1 chloroform:methanol:water ratio. Samples were vortexed, centrifuged to separate the organic solvent and aqueous phases, and the organic solvent phase was collected. Extraction was repeated a total of 3 times. The organic solvent phases were combined and washed twice with 1 ml 1.0 M KCl. Prior to TLC, lipid samples were dried under a stream of argon gas and resuspended in 1:1 chloroform:methanol to a final concentration corresponding to 4 μl of solvent per 1 mg cell pellet wet weight. Isolated lipids were spotted onto heated glass-backed silica gel 60 plates (1057210001; Millipore Sigma), and neutral lipids were separated in a mobile phase of 80:20:1 hexane:diethyl ether:glacial acetic acid. TLC bands were visualized by spraying dried plates with cupric acetate in 8% phosphoric acid and baking at 145°C for an hour.

### TLC quantification

Stained TLC plates were scanned and then processed for quantification using Fiji (ImageJ). Each plate was spotted with a neutral lipid reference standard mixture (Cat # 18-5C; Nu-Chek Prep). The standard was prepared in chloroform to a final concentration of 10 mg/ml and diluted to 1µg/µL before loading onto plate. The neutral lipid standard was used to create a standard curve in which the x-axis displayed the calculated lipid mass in micrograms, and the y-axis displayed the band intensity estimated by using Fiji.

### Yeast LD number and area quantification

For Fig 7B TEM images, LDs were counted by hand using the Fiji multipoint tool. The area of these same LDs was determined by tracing the perimeter of each by hand using the Fiji freehand line tool. Each LD was selected as an ROI, then the area quantified using the “Measure” tool in Fiji and reported in µm^2^. For fluorescence images in Fig 9A, LD number per cell was quantified by counting MDH-stained LDs, by hand, using the Fiji multipoint tool.

### Mammalian LD area quantification

For Fig 7G images, LD area was quantified using the Trainable WEKA Segmentation plugin from FIJI. Using max-projections, ROIs were drawn around the boundaries of each individual cell to measure the cell area. A WEKA classifier (Arganda-Carreras et al., 2017) was trained to segment MDH-stained LDs from cellular background. After thresholding the classified images, particles were analyzed for each ROI and the LD area per cell was recorded. The LD area per total cell area was calculated and analyzed in GraphPad Prism 8.

### Fluorescent signal quantification of Tld1 and Tgl3 imaging under cerulenin treatment

In Fig 8F, fluorescent signals for Tld1-GFP and Tgl3-mRuby foci were quantified from confocal maximal projections from Fig 8E imaging using Fiji. To summarize, for each image, the midplane z-section of the DAPI channel (MDH-stained LDs) was converted to grayscale, then random LDs were selected using the oval selection tool. Each of these LDs were marked as individual ROIs, along with a random area with no fluorescent signal selected as background, then all were saved to the ROI manager. Next, the maximal projections for the DAPI, RFP, and GFP channels were merged into one image, and the previously selected LD ROIs were overlaid on to image. The fluorescent signal for each channel, represented as Raw Integrated Density, was then measured for each ROI. These values were then subtracted from the background ROI integrated density for each channel to obtain a Tld1-GFP and Tgl3-mRuby signal value for each ROI. Then, for both GFP and mRuby channels, each ROI signal measurement was divided by the ROI with highest Raw Integrated Density to obtain a ratio (Raw Integrated Density / Max Integrated Density). For each ROI, said ratio for Tld1-GFP signal and Tgl3-mRuby signal were plotted against each other for “No Cerulenin” and “3hr Cerulenin” conditions. Pearson’s correlation coefficient (r) was calculated for both graphs.

### Statistical analysis

Graphpad Prism 8 software was used to perform all statistical analyses, with graphs indicating the mean + standard deviation. Two-tailed, unpaired t tests were performed with Welch’s correction. Where indicated, ordinary one-way ANOVA tests were performed, with Tukey’s multiple comparisons test applied. For both t tests and ANOVA, ns, P ≥ 0.05; *, P < 0.05; **, P < 0.01; ***, P < 0.001; ****, P < 0.0001.

### Conventional TEM

Yeast cells were grown in the desired conditions and processed in the University of Texas Southwestern Electron Microscopy Core Facility using a adapted protocol from Wright (Wright, 2000). In brief, cells were fixed in potassium permanganate, dehydrated, and stained in uranyl acetate and embedded in Spurr Resin. Specimen blocks were polymerized at 60°C overnight and sectioned at 70 nm with a diamond knife (Diatome) on a Leica Ultracut UCT 6 ultramicrotome (Leica Microsystems). Sections were poststained with 2% uranyl acetate in water and lead citrate. Sections were placed on copper grids (Thermo Fisher Scientific). Images were acquired on a Tecnai G2 spirit TEM (FEI) equipped with a LaB6 source at 120 kV by using a Gatan Ultrascan charge-coupled device camera.

### Whole cell protein extraction and sample preparation

Whole cell protein extracts were isolated from 25 OD_600_ units of cells. Pellet wet weights were normalized prior to freezing at −20°C. Frozen cell pellets were incubated with 20% trichloroacetic acid (TCA) for 30 min on ice with occasional mixing using a vortex. Precipitated proteins were pelleted in a 4°C centrifuge at 16,000 g for 5 min. After removing the supernatant, the pellet was washed three times with cold 100% acetone followed by brief sonication. After the washes, the protein pellets were dried in an RT speed vac for 15 min to remove residual acetone. Dried protein pellets were neutralized with 1.5M Tris-HCl pH 8.8, then resuspended directly in 250µL of 1X Laemelli sample buffer (Laemmli, 1970). Samples were briefly sonicated and boiled at 95°C for 5 min. Fig 1D, 1F, and 7F protein samples were extracted as described above, except following neutralization, protein pellets were resuspended in 250µL resuspension buffer (50mM Tris pH 6.8, 1mM EDTA, 1% SDS; 6M Urea, 1X Halt Protease and Phosphatase Inhibitor Cocktail [78441; Thermo Fisher Scientific], and 1% beta-mercaptoethanol). These samples were sonicated briefly, but not subjected to heating/boiling to prevent aggregation of these hydrophobic droplet proteins. 2X Laemelli sample buffer was added to these samples immediately prior to gel loading.

### Immunoblot analysis

Following protein extraction, samples were pelleted at 16,000 g for 3 min to remove insoluble debris. Equal volumes of each sample were then subjected to SDS-PAGE and western blot analysis. Proteins were separated on a precast Mini-PROTEAN® TGX™ 10% SDS-PAGE gel (4561034; BioRad) and then transferred to a 0.45 μm nitrocellulose membrane in Towbin SDS transfer buffer (25 mM Tris, 192 mM glycine, 20% methanol, and 0.05% SDS; pH 8.2) using a Criterion tank blotter with plate electrodes (1704070; BioRad) set to 70V constant, for 1hr. Immediately after transfer, membranes were stained with PonceauS, imaged on a ChemiDoc™ Touch Gel Imager (1708370; BioRad) and cut using a clean razor blade. Membranes were blocked with 5% milk dissolved in Tris-buffered saline +Tween (TBS-T) buffer, and primary antibodies were allowed to bind overnight at 4°C. Primary antibodies used for determining protein expression are as follows: GFP (ab290; 1:5,000 dilution; Abcam), GAPDH (ab9485; 1:2,500 dilution; Abcam), mNeonGreen (Cat# 32f6; 1:1,000 dilution; ChromoTek). Immunoblots were developed by binding HRP-conjugated anti-rabbit IgG (ab6721;1:5,000; Abcam) or anti-mouse IgG (ab6728; 1:1,000; Abcam) secondary antibodies to the membrane for 1 h in the presence of 5% milk followed by four washes in TBS-T and developing with ECL substrate (1705061; BioRad). Blot signal was captured using the same BioRad ChemiDoc™ Touch Gel Imager, as noted above. Protein expression levels were quantified by measuring band intensity using ImageJ and normalizing these values to wildtype to generate an abundance value relative to control.

### Whole cell protein extraction for immunoprecipitation

Yeast were collected and prepared as described above. The samples were subjected to a modified cold glass bead cell lysis and protein extraction protocol (DeCaprio and Kohl, 2020). In brief, cells were washed in cold tris-buffered saline and pelleted at 2000 g for 5 min at 4°C. Yeast pellets were resuspended in ice-cold lysis buffer plus protease inhibitors (50mM Tris-HCl pH 7.5, 120mM KCl, 5mM EDTA, 0.1% Nonidet P-40 Substitute, 10% Glycerol, 1mM DTT, 1mM PMSF, and 1X Halt Protease and Phosphatase Inhibitor Cocktail [78441; Thermo Fisher Scientific]), transferred to a 2 mL screw-cap microcentrifuge tube (Cat # 02-681-343; Fisher Scientific) containing glass beads (Cat # G8772-500G; Milipore Sigma), and lysed 3 times in a MiniBeadBeater for 90 sec each at 4°C. In between bead beating, samples were chilled in an ice bath for 2 min. Samples were then pelleted at 1000 g at 4°C for 30 sec. Supernatants were transferred to a 1.5 mL microcentrifuge tube, and beads in screw-cap tubes were washed once again in the same lysis buffer plus protease inhibitors and pelleted like above. Supernatants of screw-cap tubes were transferred to same 1.5 mL tube as above, and were cleared of insoluble debris, twice at 16000 g for 10 min at 4°C. A final clearance spin of lysates was done at 20000 g for 30 min at 4°C. Protein concentrations were then quantified using the Pierce™ BCA Protein Assay Kit (Cat # 23227; Thermo Fisher Scientific) in a 96-well plate format (Cat # 353072; Corning). Sample absorbances were measured at 562 nm using a VersaMax Microplate Reader and SoftMax Pro Software. Absorbances were converted to protein concentration using a bovine serum albumin standard curve.

### Immunoprecipitation (IP)

For immunoprecipitation, an mNeonGreen-Trap Agarose Kit (ntak-20; Chromotek) to pull down Tld1-mNeonGreen (mNG) fusion protein was used, according to manufacturer’s protocol. To begin, for each sample 25µL of agarose beads containing an anti-mNG nanobody were washed in 500µL of ice cold dilution buffer (10mM Tris-HCl pH 7.5, 150mM NaCl, 0.5mM EDTA, and 0.018% sodium azide), centrifuged down at 2500 g for 5 min at 4°C, and buffer removed. 4000 µg of protein lysate from cold glass bead lysis for each sample was centrifuged at 16000 g, 5 min, at 4°C. Then, lysates were incubated with the washed mNG beads and rotated end over end for 1 hr at 4°C. Samples were then spun down at 2500 g for 5 min at 4°C and supernatants removed. Beads were then washed three times in 500µL wash buffer (10 mM Tris/Cl pH 7.5, 150 mM NaCl, 0.05 % Nonidet™ P40 Substitute, 0.5 mM EDTA, and 0.018 % sodium azide) and centrifuged like above, in between each wash. After final wash and spin, supernatant was removed, beads were transferred to a fresh 1.5 mL tube. 2X Laemmli sample buffer was added to beads, and samples were boiled for 5 min at 95°C

### LC-MS/MS proteomics

Following boiling step, IP samples were centrifuged at 2500 g, for 2 min at 4°C to pellet beads. The entirety of each supernatant was loaded onto a 10% mini-protean TGX gel (4561033; Bio-Rad). Samples were subjected to electrophoresis at 90 V constant until the dye front was ∼10 cm into the gel. The gel was subsequently removed from the casing and stained with Coomassie reagent (0.5 Coomassie G-250, 50% methanol, 10% acetic acid) for 10 min on an RT rocker. The gel was then rinsed three times in sterile Mili-Q water to gently destain. Once the gel was sufficiently destained, 10-cm gel bands were excised from each lane, taking care to exclude the stacking gel and dye front. Gel bands were further cut into 1-mm squares and placed into sterile microcentrifuge tubes. Samples were digested overnight with trypsin (Pierce) following reduction and alkylation with DTT and iodoacetamide (Sigma-Aldrich). The samples then underwent solid-phase extraction cleanup with an Oasis HLB plate (Waters), and the resulting samples were injected onto an Orbitrap Fusion Lumos mass spectrometer coupled to an Ultimate 3000 RSLC-Nano liquid chromatography system. Samples were injected onto a 75 μm i.d., 75-cm long EasySpray column (Thermo Fisher Scientific) and eluted with a gradient from 0 to 28% buffer B over 90 min. The buffer contained 2% (vol/vol) acetonitrile and 0.1% formic acid in water, and buffer B contained 80% (vol/vol) acetonitrile, 10% (vol/vol) trifluoroethanol, and 0.1% formic acid in water. The mass spectrometer operated in positive ion mode with a source voltage of 1.5–2.0 kV and an ion transfer tube temperature of 275°C. MS scans were acquired at 120,000 resolution in the Orbitrap, and up to 10 MS/MS spectra were obtained in the ion trap for each full spectrum acquired using higher-energy collisional dissociation for ions with charges 2–7. Dynamic exclusion was set for 25 s after an ion was selected for fragmentation. RawMS data files were analyzed using Proteome Discoverer v 2.4 (Thermo Fisher Scientific), with peptide identification performed using Sequest HT searching against the *Saccharomyces cerevisiae* protein database from UniProt. Fragment and precursor tolerances of 10 ppm and 0.6 dalton were specified, and three missed cleavages were allowed. Carbamidomethylation of Cys was set as a fixed modification, with oxidation of Met set as a variable modification. The false-discovery rate cutoff was 1% for all peptides.

### Cell culture

U2-OS cells and Hela cells were cultured in DMEM (D5796; Sigma) supplemented with 10% Fetal Bovine Serum (F4135; Sigma), 1% penicillin streptomycin solution (30-002-Cl; Corning), and 25mM HEPES (H0887; Sigma-Aldrich). The cells were passaged when they reached 80–90% confluence with 0.25% trypsin-EDTA (25-053-Cl; Corning). To promote TG-rich LD biogenesis, cells were incubated with 600 μM of OA conjugated with 100 μM of FA-free BSA (A8806; Sigma-Aldrich) for 16 hours. To promote SE-rich LD biogenesis, cells were incubated with 200 μM of cholesterol conjugated with methyl-β-cyclodextrin (C4555; Sigma-Aldrich) for 16 hours. OA and CHO were added to DMEM supplemented with 25mM HEPES.

### Cloning and transient transfection

Full length Tld1-EGFP and fragments were generated after PCR amplification of Tld1 from yeast pBP73G Tld1 untagged overexpression plasmid and cloning into pEGFP-N2 (XhoI/BamHI). Tld1-EGFP point mutants were generated by overlap extension PCR amplification of full length Tld1 from yeast genomic DNA. pEGFP-N2 alone, served as a negative control. The plasmids were transfected into U2-OS and HeLa cells using Lipofectamine 3000 Transfection Reagent (L3000001; Invitrogen) and Opti-MEM (31985-070; Gibco) for 48 h before experiments.

### IF staining

Control and OA-treated transfected cells were fixed with 4% PFA solution in PBS for 15 min at RT. For IF staining, the cells were washed with PBS, permeabilized with 0.5% Triton X-100 (X100; Sigma-Aldrich) in PBS at RT for 5 min. Cells were washed and blocked in IF buffer (PBS containing 3% BSA) for 30 min. Cells were then incubated with primary antibody in IF buffer for 1 h, washed thrice with PBS, incubated with secondary antibody in IF buffer for 1 h, and given two washes with PBS. Control and OA-treated transfected cells were then incubated with MDH AutoDOT (SM1000a; 1:1,000 dilution; Abcepta) for 20 min, washed thrice with PBS, and then stored in PBS at 4°C before imaging. The primary antibody used was mouse anti-Hsp90B1 (AMAb91019; 1:100 dilution; Sigma-Aldrich). The secondary antibody used was donkey anti-mouse Rhodamine Red-X (715-295-151; 1:1,000 dilution; Jackson Laboratories). CHO-treated Hela cells were fixed with 4% formaldehyde in PBS for 15 min at RT. Transfected cells were washed thrice with PBS and stored in PBS at 4°C before imaging. Non-transfected control cells were permeabilized with 0.5% Triton X-100 in PBS at RT for 5 min. Cells were washed thrice with PBS, incubated with 2uM BODIPY 493/503 (D3922; Invitrogen) in PBS at RT for 15 min, given two washes with PBS, and stored in PBS at 4°C before imaging. TG-rich LDs were visualized by staining the cells with AutoDOT, and SE-rich LDs were visualized by staining with BODIPY 493/503.

### Fluorescence microscopy

For confocal microscopy, yeast cells were grown as described above and collected by centrifugation at 4,000 rpm for 5 min. Where indicated, cells were incubated for 5 min with MDH (SM1000a; Abcepta) at a final concentration of 0.1 mM to visualize LDs. Before imaging, yeast cells were washed with 1 ml of Mili-Q water and resuspended in 50-100µL of Mili-Q water. Mammalian cells were imaged in 8-well Nunc™ Lab-Tek™ II chambered coverglass (Cat #154409; Thermo Scientific). All images were taken as single slices at approximately mid-plane using a Zeiss LSM880 inverted laser scanning confocal microscope equipped with Zen software. Images were taken with a 63x oil objective NA = 1.4 or 40x oil objective NA = 1.4 at RT, unless noted otherwise. Approximately seven Z-sections of each image were taken for yeast, and five for mammalian cells. Merged images were maximum intensity z-projections (generated by Fiji), unless indicated otherwise in Figure legends. For epifluorescence microscopy, cells were grown, stained, and collected as described above. Imaging was performed on an EVOS FL Cell Imaging System at RT.

### Polarized light microscopy

To detect smectic LC phase in SE-rich LDs HeLa cells were imaged using polarized light microscopy. All images were taken as 31 z-sections using Nikon eclipse Ti inverted confocal microscope equipped with built-in polarizer and analyzer filters, and Ni elements software. Images were taken with a 100x oil objective NA=1.45 at RT.

### Yeast strain generation and plasmid construction

A modified version of the lithium acetate method was used for the generation of all yeast knock outs and knock ins. Briefly, yeast were diluted from a ∼24h culture to an OD_600_ = 0.001 in YPD media and allowed to grow 16-20h, overnight, until they reached OD_600_ = 0.6. For each transformation, the entire culture was pelleted (50mL), washed with sterile Mili-Q water, washed with 0.1 M lithium acetate, pelleted and resuspended in 1mL 0.1M lithium acetate. 100µL this yeast-lithium acetate suspension was added to ∼1mL of transformation solution (40% polyethylene glycol in 0.1 M lithium acetate, 0.25 μg/μl single-stranded carrier DNA [D9156; Sigma-Aldrich]) supplemented with 5-10µg of PCR product. Transformations were vortexed and incubated at 30°C for 45 min, then 42°C for 30 min. Cells were then pelleted at 2000 g, 2 min and gently washed with sterile Mili-Q water, then pelleted again. For antibiotic marker transformations, yeast were then resuspended in 2mL fresh YPD media and allowed to recover overnight, 30°C, 225 RPM. The following day, cells were pelleted and plated onto YPD plates containing antibiotic and incubated at 30°C, 2-3 d. For auxotrophic marker transformations, yeast were plated onto SC dropout plates same day (immediately after Mili-Q washing step) and incubated at 30°C, 2-3 d. Plasmids were generated for this study using Gibson Assembly following the manufacturer’s protocol (E2611; NEB). All pBP73-G vectors were cut with XbaI and XhoI. All pRS305 vectors were cut with SacI and XhoI. For yeast plasmid transformations, cells were grown in YPD media, overnight until saturation. 1 mL of overnight culture was pelleted at 12000 RPM, 2 min at RT. Pellets were then washed in 0.1M Lithium Acetate and centrifuged again, like above. Yeast cells were then resuspended in ∼300 µL transformation solution (40% polyethylene glycol in 0.1 M lithium acetate, 0.25 μg/μl single-stranded carrier DNA [D9156; Sigma-Aldrich]) with 1µg of plasmid DNA, vortexed briefly, and incubated at RT for 1hr. Transformations were then gently mixed, DMSO added to a final concentration of 10%, and heat shocked at 42°C, 10 min. Samples were then put on ice for 2 min, then entire reaction was plated onto SCD plates lacking uracil or leucine, and incubated at 30°C for 2-3 days.

### Proteomics quantification

Proteomics quantification and analysis were performed using Excel. All samples were analyzed in triplicate. To adjust for total protein differences between samples, the sum of all spectral counts within each sample was taken and divided by the average of the spectral count sums in the empty vector soluble mNG (EV-mNG) samples. This ensured differences observed in the proteomics data are not due to unequal “loading” into the MS. Next, only proteins with detectable spectral counts in all 3 replicates of the Tld1-mNG IP samples were considered for analyses, regardless of whether they were present in the EV-mNG IP replicates. From this list proteins, those with undetectable spectral counts in the EV-mNG IP replicates had their spectral counts changed from “0” to “1” to aid in quantifications for statistical analysis. To generate a high-confidence list of Tld1 interacting proteins, the average spectral counts of each protein from the Tld1-mNG IPs were divided by the corresponding average spectral counts from the EV-mNG IP samples. Therefore, proteins more abundant in the Tld1-mNG IP samples would produce a ratio >0. To generate volcano plots in GraphPad Prism, log_2_ values were calculated for the ratio of average protein expression in EV-mNG and Tld1-mNG (i.e., log_2_[protein A in Tld1-mNG/protein A in EV-mNG]). Then, the p-value for significance of the abundance for each protein in EV-mNG and Tld1-mNG replicate samples was calculated via t test. Finally, the -log_10_ of these p-values was calculated and plotted against the above log2 values in volcano plot form. Significance cut-off on the y-axis was the -log10 of P = 0.05, or 1.3.

### Cartoon development

All cartoons created with BioRender.com (Figure 9C) or Microsoft Powerpoint (all others). For Fig 2A, the hydrophobicity plot was generated using data collected from Phobius open access hydrophobicity predictor (Käll et al., 2004) and the helical wheel generated using HeliQuest (Gautier et al., 2008).

## Figure Supplements

**Figure Supplement 1:**
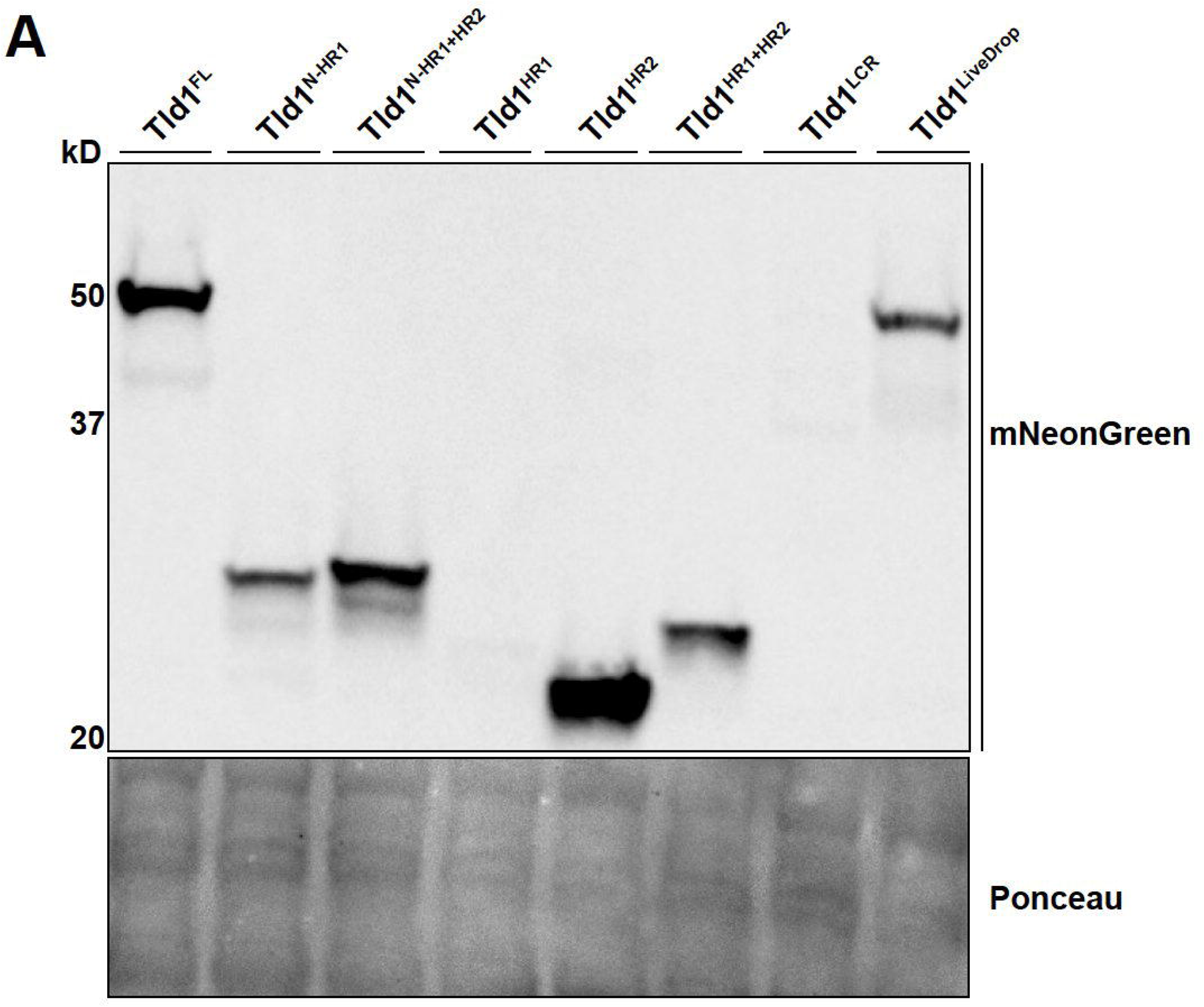
Whole cell protein expression levels of Tld1 fragments. **(A)** Western blot of yeast strains overexpressing Tld1 fragments, tagged with a mNeonGreen fluorophore. Membranes were blotted with anti-mNeonGreen antibody and Ponceau S stain served as loading control for total protein. LCR=Low Complexity Region.

**Figure Supplement 2:**
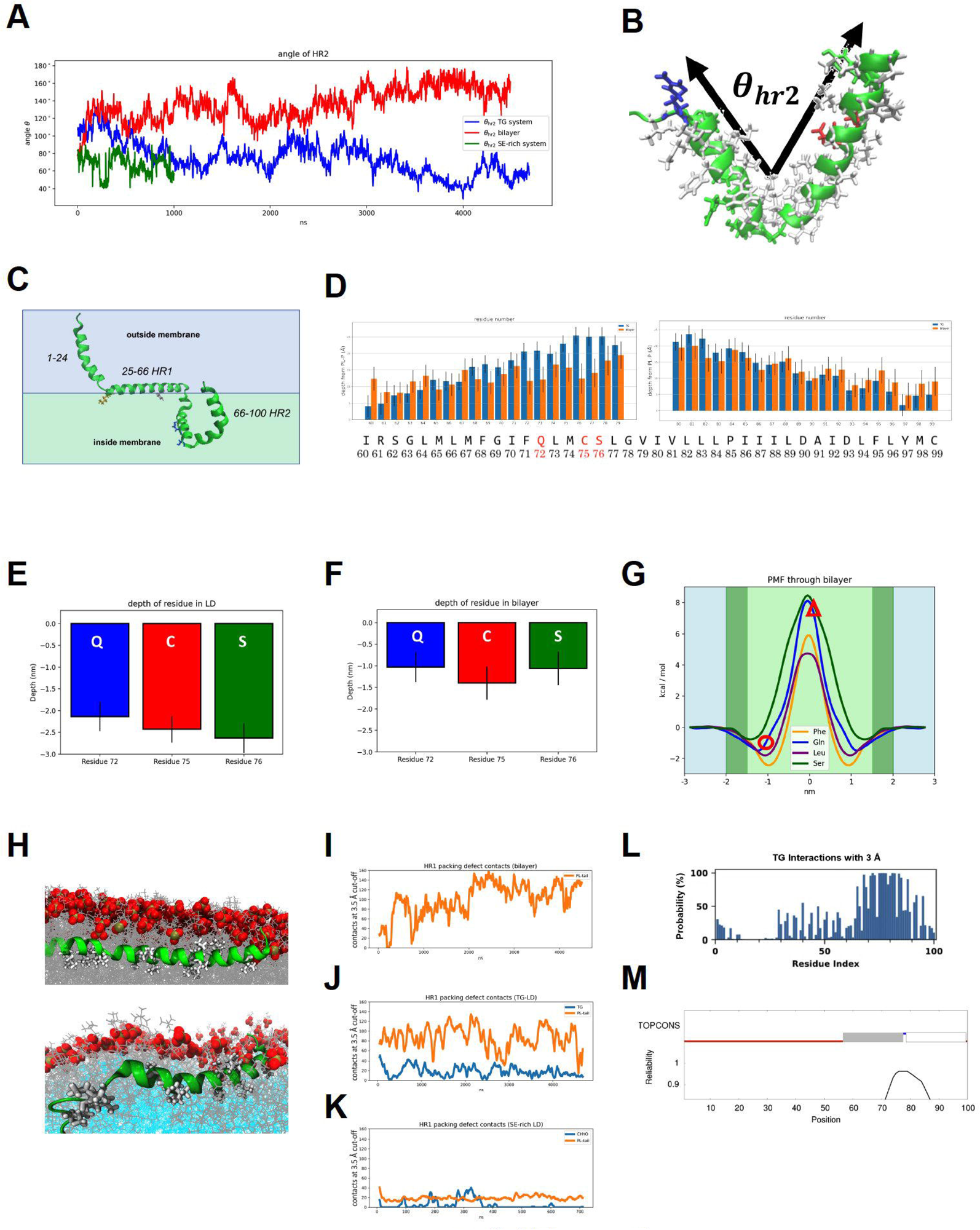
MD analysis of Tld1 HR1 and HR2 conformations. **(A)** Angle of HR2 over time. **(B)** The angle was defined by the two endpoints (residues 61 and 100) and kink (residue 78). The predicted/initial angle was 100 degrees. **(C)** The predicted structure of Tld1^N+HR1+HR2^ through RoseTTAFold. Gln-72 and Ser-76 are blue, Phe-44 is pink, and Lys-26 is orange. **(D)** Average depths from of residues 60-99 (sidechains) below the PL phosphorous plane in the TG-rich LD and ER bilayer. Focusing on the polar residues, the average depth of the residue’s COM is significantly deeper in the TG-LD **(E)** than in the ER bilayer **(F)**. **(G)** Free energy profiles for membrane permeation show stability of GLN and SER ∼1nm below the phosphate plane just under the headgroups (dark green regions) and unfavorable penalty for pulling them ∼2 nm below the plane into the PL tail region (light green region). The depths of Gln72 and Ser76 are marked by a red circle for the ER bilayer and a triangle for the LDs. **(H)** HR1 sequence interacting with the bilayer (top) and TG-LD (bottom). **(I)** In the ER bilayer, these contacts are all PL-tail interactions. **(J)** In the TG-LD system, there is a combination of PL-tail and TG defects interactions. **(K)** In the 90:10 CHYO:TG LD (SE-rich LD), the interactions rarely occur as there are too few packing defects. **(L)** The probability (y-axis) of each residue (x-axis) interacting with a TG molecule. HR2 is in consistent contact with TG molecules. **(M)** TOPCONS result of predicting transmembrane sequences. The grey and white bar represent transmembrane sequences, which correspond to the HR2 helices.

**Figure Supplement 3:**
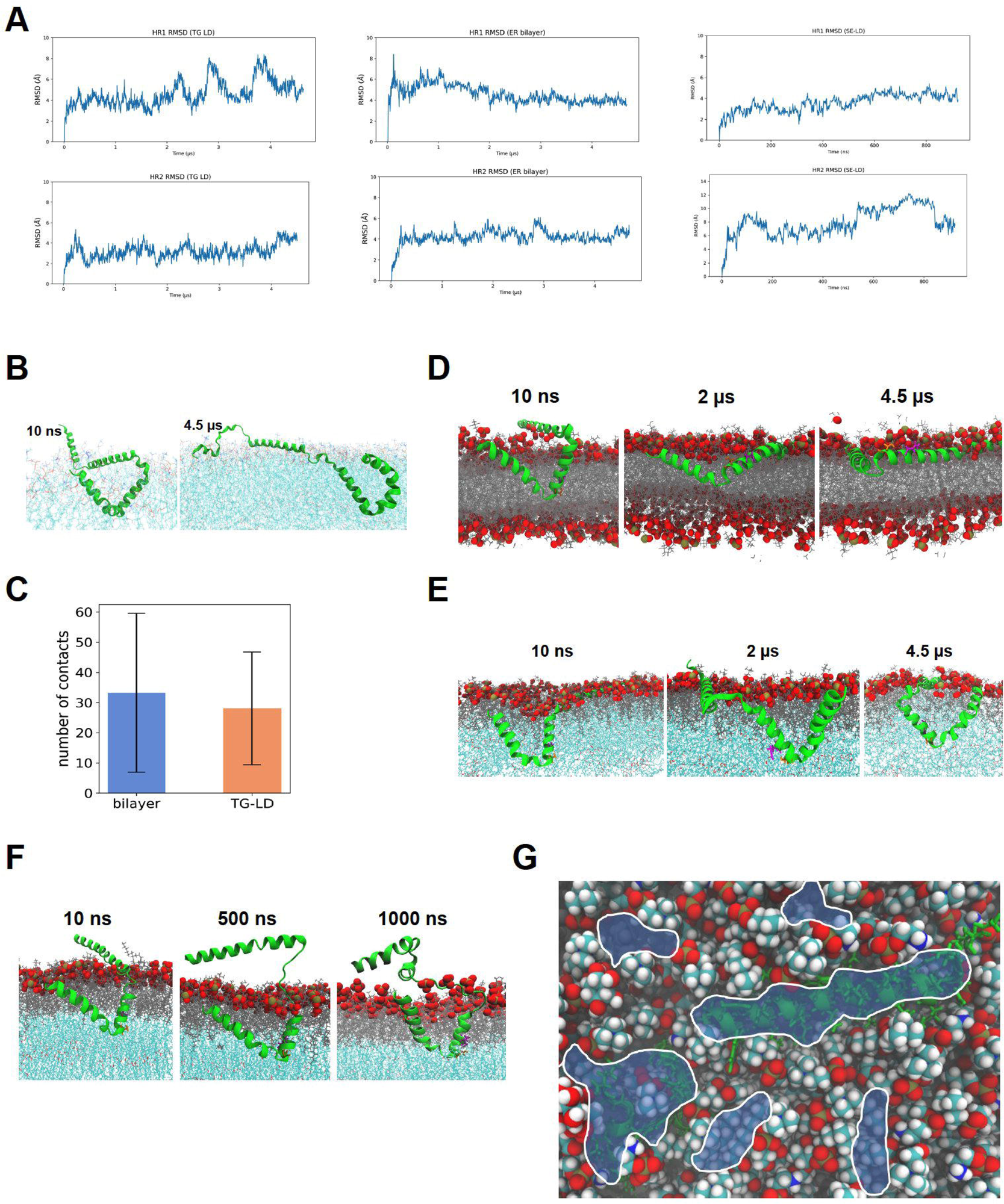
Tld1 HR1 and HR2 conformational changes in LDs versus the ER. **(A)** RMSD of HR domains in TG (left) and SE-rich LDs (right) and ER bilayer (middle). **(B)** Tld1^N-HR1+HR2^ at the beginning and end of the simulation of the TG-LD system. The N terminus partially unfolds and makes some interactions with PL headgroups. **(C)** The nature and number of interactions for the bilayer and TG-LD remain similar throughout the trajectories. **(D)** Evolution of Tld1^HR1+HR2^ in ER bilayer. Binding of HR1 occurs within the first 50 ns, while the kink with the polar residues rises to the surface. **(E)** Evolution of Tld1^HR1+HR2^ in TG-rich LD. Binding of HR1 occurs withing the first 50 ns, while the kink remains in the LD core. **(F)** Evolution of Tld1^HR1+HR2^ in SE-rich LD. Binding of HR1 never occurs, but kink remains in the LD core. **(G)** Packing defects (shaded blue) over HR1 (right) and HR2 (left) in TG-LD.

**Figure Supplement 4:**
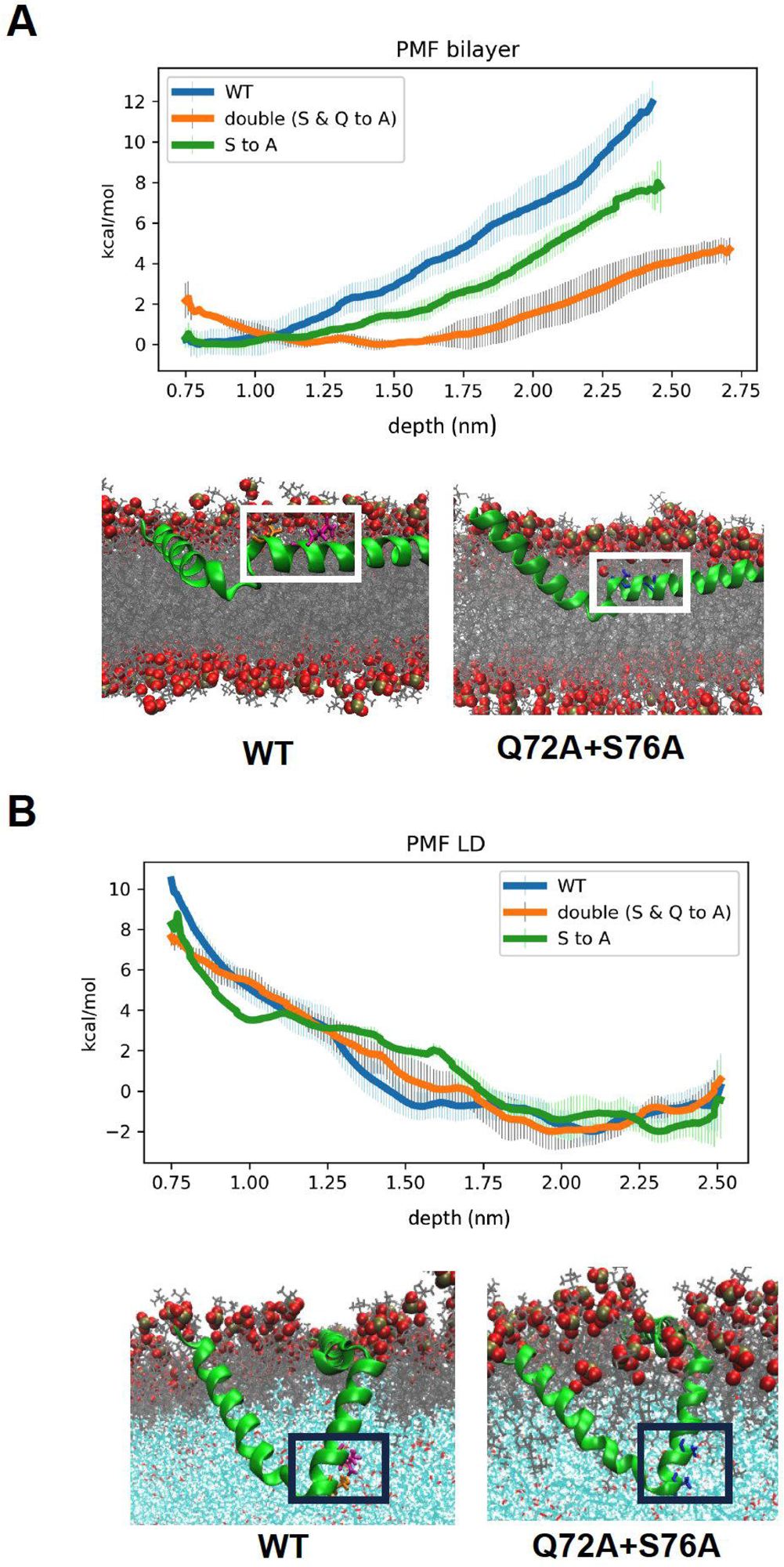
MD analysis of Tld1 HR2 point mutants in the LD monolayer versus ER bilayer. **(A,B)** Free energy profiles for kink formation in the ER bilayer (A) and TG-rich LD (B). The x-axis is the depth of the kink residues below the PL phosphate plane. Representative conformations for free energy minima show the Q72A+S76A mutant has its kink region deeper in the ER membrane but remains in the same position as the WT in the LD.

**Figure Supplement 5:**
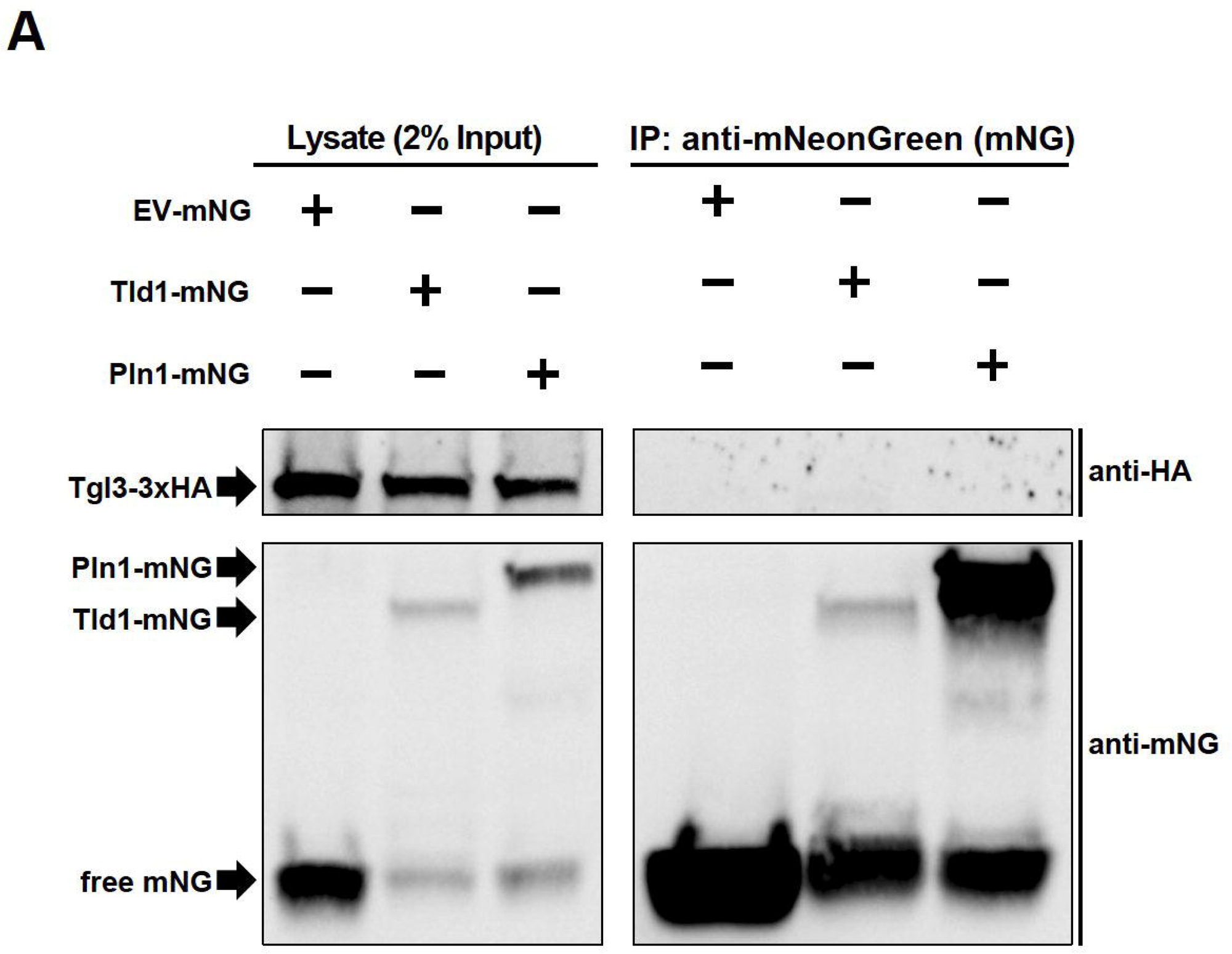
Co-immunoprecipitation (Co-IP) and western blot of Tld1. **(A)** Co-IP and western blot of yeast strains endogenously tagged with Tgl3-3xHA and overexpressing either Tld1-mNeonGreen (Tld1-mNG), Pln1-mNG, or empty vector-mNG (EV-mNG). Membranes were blotted with anti-mNG and anti-HA antibodies. Representative of three independent experiments.

